# Structural insights into the mammalian late-stage initiation complexes

**DOI:** 10.1101/779504

**Authors:** Angelita Simonetti, Ewelina Guca, Anthony Bochler, Lauriane Kuhn, Yaser Hashem

**Affiliations:** Université de Strasbourg, CNRS, Architecture et Réactivité de l’ARN, UPR9002, Strasbourg 67000, France; INSERM U1212 Acides nucléiques : Régulations Naturelle et Artificielle (ARNA), Institut Européen de Chimie et Biologie, Université de Bordeaux, Pessac 33607, France; Proteomic Platform Strasbourg - Esplanade, Institut de Biologie Moléculaire et Cellulaire, CNRS, Université de Strasbourg, Strasbourg 67000, France

**Author notes:** These authors contributed equally (Angelita Simonetti, Ewelina Guca).

## Abstract

In higher eukaryotes, the mRNA sequence in direct vicinity of the start codon, called the Kozak sequence (CRCCaugG, where R is a purine), is known to influence the rate of the initiation process. However, the molecular basis underlying its role remains poorly understood. Here, we present the cryo-electron microscopy (cryo-EM) structures of mammalian late-stage 48S initiation complexes (LS48S IC) in the presence of two different native mRNA sequences, β-globin and histone 4 (H4) at overall resolution of 3Å and 3.5Å, respectively. Our high-resolution structures unravel key interactions from the mRNA to eukaryotic initiation factors (eIF): 1A, 2, 3, 18S rRNA, and several 40S ribosomal proteins. In addition, we were able to study the structural role of ABCE1 in the formation of native 48S ICs. Our results reveal a comprehensive map of the ribosome/eIFs –mRNA and –tRNA interactions and suggest the impact of mRNA sequence on the structure of the LS48S IC.

## INTRODUCTION

mRNA translation initiation in mammals is more complex than its bacterial counterpart. Indeed it includes more steps, more initiation factors and more regulation pathways. One can summarize the overall process in four steps, starting with pre-initiation. During pre-initiation, the ternary complex (TC) is formed by the binding of the heterotrimeric eukaryotic initiation factor 2 (eIF2) to one molecule of guanosine triphosphate (GTP) and the initiator methionylated tRNA (tRNA_i_^Met^). The TC then binds to the post-recycled ribosomal small subunit (SSU), also called 40S subunit. TC recruitment is partially mediated by eukaryotic initiation factors attached to the 40S, eIF1, eIF1A, and 13-subunit eIF3 complex. This leads to the formation of the 43S pre-initiation complex (PIC). The architecture of the 43S PIC has been investigated structurally at low to intermediate resolutions (Aylett et al., 2015; Erzberger et al., 2014; Hashem et al., 2013).

The second step consists on the recruitment of the 5’ capped mRNA and leads to the formation of the 48S IC. This step is mediated by the cap-binding complex composed of eIF4F, eIF4A and eIF4B (Gross et al., 2003; Jackson et al., 2010; Marintchev et al., 2009; Rogers et al., 2001).

The third step is the scanning process for the start codon (AUG) in the 5’ to 3’ direction. This step was first investigated structurally in yeast (Hussain et al., 2014; Llacer et al., 2015) using *in vitro* reconstituted complexes. Upon start-codon recognition, the codon: anticodon duplex is formed between the mRNA and the tRNA_i_^Met^ aided by the eIF1A N-terminal tail (NTT) (Hinnebusch, 2011; Llácer et al., 2018; Lomakin and Steitz, 2013). The GTP is hydrolysed by eIF2γ, eIF1 dissociates from the P-site along with eIF1A C-terminal tail (Zhang et al., 2015) and the N-terminal domain (NTD) of eIF5 takes their place on the 40S (Llácer et al., 2018), before eIF5-NTD dissociates in turn at a stage that still remains to be elucidated. This results in the formation of the LS48S IC (that we describe in this work). The arrest of scanning and a cascade of structural rearrangements lead to the sequential dissociation of most eIFs upon the release of inorganic phosphate (Pi), generated from the GTP hydrolysis. eIF3 stays attached to the remaining complex probably through its peripheral subunits and leaves at a later stage during early elongation cycles (Beznosková et al., 2013, 2015). During all these steps the post-recycling factor ABCE1 can bind directly to the 40S and act as an anti-ribosomal subunits association factor (Heuer et al., 2017; Kiosze-Becker et al., 2016; Mancera-Martínez et al., 2017).

In the final fourth step, ABCE1 is replaced by the GTPase eIF5B on helix 14 of 18S rRNA, thus stimulating the joining of the 40S and 60S ribosomal subunits, forming an 80S complex (Fernandez et al., 2013) and eIF1A and eIF5B are released together (Fringer et al., 2007).

The sequences flanking the AUG start-codon region have been identified as crucial for start site selection by the IC (Kozak, 1986, 1987b, 1987a, 1989). The optimal sequence for translation initiation in eukaryotes was named after Marylin Kozak, who first defined the optimal sequence in vertebrates as CRCCaugG, where R stands for a purine (Kozak, 1984, 1989). In this motif, modification of certain positions have influence on translation efficiency, such as (−3) and (+4) (Kozak, 1984). As a result, a sequence can be dubbed “strong” or “weak” by considering those positions. It was further shown that the substitution of A(−3) for pyrimidine, or mutations of the highly conserved G(+4), lead to a process known as “leaky scanning” with bypass of the first AUG and initiation of translation at the downstream start codon (Kozak, 1986, 1989; Lin et al., 1993). More recent studies observed a more extreme case of sequence-dependent translation initiation regulation, dubbed “cap-assisted” for certain cellular mRNAs, such as those encoding histone proteins and in particular histone 4 (H4) mRNA (Martin et al., 2011, 2016). Cap-assisted internal initiation of H4 mRNA implies a very minimalistic scanning mechanism, which is possible thanks to the presence of a tertiary structure on the mRNA at the channel entrance. This element assists in placing the start codon very close to the P-site almost immediately upon its recruitment through the cap-binding complex.

In spite of the tremendous recent advances in understanding this phase of translation, high-resolution structural studies of the initiation process have been conducted by *in vitro* reconstitution of the related complexes. This approach often requires biologically irrelevant molar ratios of the studied eIFs (Aylett et al., 2015; Erzberger et al., 2014; Hashem et al., 2013; Hussain et al., 2014; Llacer et al., 2015), thus limiting insight into more subtle regulatory pathways. Moreover, the structures of the mammalian (pre)ICs are still at intermediate resolutions approximating 6Å (Eliseev et al., 2018; des Georges et al., 2015; Simonetti et al., 2016). Finally, although the role of ABCE1 was experimentally demonstrated as a ribosomal subunit anti-association factor preventing premature binding of the 60S (Heuer et al., 2017), its impact on the 48S complex formation and conformation is still unclear.

Here, we present cryo-EM structures of LS48S IC formed after recognition of the start codon on two different native and abundant cellular mRNAs, β-globin and histone 4, presenting variants of the Kozak sequence. Both complexes were prepared and isolated in near-native conditions from rabbit reticulocyte lysate (Figure 1A). Although the initiation regulation may differ mechanistically between these two archetype mRNAs, our structures provide a high-resolution snapshot on the Kozak sequence-dependent variable interactions in the LS48S IC in mammals.

**Figure 1.**
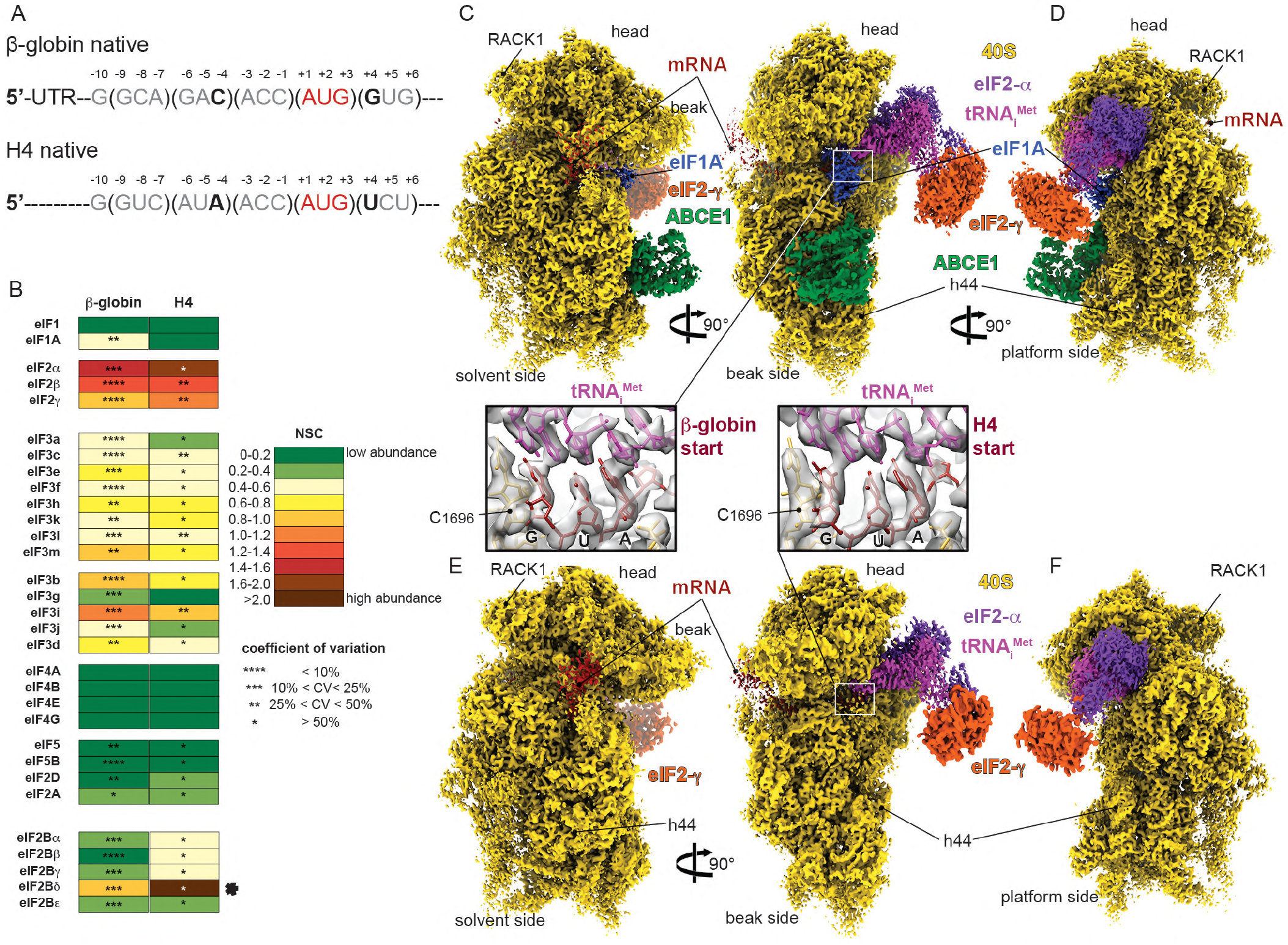
Overall structure of β-globin and H4 late-stage 48S initiation complexes. (A) mRNA sequences used to form and purify the β-globin and the H4 ICs. Only the sequences near the AUG codon are represented and main differences in the Kozak sequence are indicated in bold. (B) Semi-quantitative mass spectrometry analysis of the eIFs in both ICs, indicating the abundance of each eIF based on the spectra count normalized. The two rounds of normalization were carried out using the total number of eIFs and estimated number of trypsin cleavage sites (see Methods). The normalized spectra counts (NSC) are presented as heat maps with cold colours indicating low abundance and warm colours indicating high abundance. The higher abundance of eIF2 proteins might be due to the excess of a free TC in the sample. The black star points out a high number of NSC for eIF2Bδ which is caused by the detection of three different isoforms of this protein. Small stars indicate the values of the coefficients of variation calculated for each NSC. In the analysis, the NSC for ABCE1 is not included, as it is a factor present also in other stages of translation than initiation. (C-D) Segmented cryo-EM reconstructions of the β-globin IC seen from (C) solvent, beak and (D) platform sides, respectively. The reconstruction shows 40S (in yellow), eIF2γ (in orange), eIF2α (in purple), tRNA ^Met^ (in magenta), mRNA (in red), eIF1A (in skyblue) and ABCE1 (in green). (E-F) Same as C-D but for the H4 IC. Boxed blowups represent the codon:anticodon duplexes in all shown reconstructions with their respective atomic models fitted in the corresponding electron densities.

## RESULTS

### Overall structure of the mammalian 48S initiation complex

The complexes were prepared using a modified version of our approach (Simonetti et al., 2016) (see Methods) that consists on stalling the LS48S IC in rabbit reticulocytes lysate (RRL) using GMP-PNP (a non-hydrolysable analogue of GTP) on the two target cellular mRNAs: mouse histone H4 mRNA (suboptimal Kozak) and human β-globin mRNA (stronger Kozak) (Figure 1A). These mRNAs were transcribed and capped from BC007075 cDNA (β-globin) and X13235 cDNA (H4). The advantage of this approach is the ability to prepare ICs bound on different mRNAs of interest directly in nuclease-treated cell extract in the absence of the endogenous mRNAs, allowing for a study of regulatory aspects of the process in natural abundance levels of native eIFs at physiological molar ratios. The composition of both complexes was investigated by mass spectrometry (Figure 1B). Our analysis reveals the incorporation in both complexes of all eIFs expected to be present after the start-codon recognition (eIF1A, eIF2α, eIF2β, eIF2γ, eIF3 complex, ABCE1). As expected, extremely poor numbers of peptides and spectra for eIF1 detected in either complex, corroborating that our complexes are at a late-stage after the start-codon recognition and eIF1 dissociation.

In parallel, we have subjected our prepared complexes to structural analysis by cryo-EM. The structure of the β-globin LS48S IC (3.0Å, 29% of the total number of 40S particles, Figures 1C-1D and S1A-S1H) shows mRNA, 40S, eIF1A, TC and ABCE1. For the H4 mRNA 48S complex (3.5Å, 6.5% of the total number of particles, Figures 1E-F and S1P-W), the main reconstruction shows mRNA, 40S and the TC. We attribute the lower percentage of H4 LS48S IC formation to contamination with 60S subunits (∼30%) (Figure S1W). Interestingly, both our cryo-EM structures and MS/MS analysis show that the H4 LS48S IC displays a significant reduction in the presence of eIF1A, leaving only residual density for its presence in the cryo-EM reconstruction (Figures 1E and S2A). Similar observation can be made for ABCE1 in the H4 LS48S IC. Our reconstructions show also another class of IC with eIF3 that is described later.

### Accommodation of the start codon in the late-stage LS48S IC

In both our reconstructions the codon:anticodon duplex is clearly formed, characterizing the cognate start-codon recognition (Figures 1C,E and 2A,B). AUG codons of both mRNAs face the _(34)_CAU_(36)_ of anticodon stem-loop (ASL) tRNA_i_^Met^, within hydrogen-bonding distances (∼2.7Å). In the case of β-globin mRNA, the codon:anticodon interaction is stabilized further by the N-terminal tail (NTT) of eIF1A (Lys7 interacts with the ribose of G(+3) from mRNA, Figure 2D). The tail also interacts with the tRNA_i_^Met^ A(35) between Gly8 and Gly9. With few exceptions, this eIF1A NTT is highly conserved among eukaryotes (Figure 2D). Recent fluorescence anisotropy with yeast reconstituted PICs (Llácer et al., 2018) demonstrated that eIF1A binds with lesser affinity to a near-cognate start codon (UUG) compared to a cognate AUG. Along the same lines, only very residual density for eIF1A can be observed in the H4 LS48S IC structure (Figure S2A) (discussed below), which reflects its weaker binding affinity after the start-codon recognition at this late stage to the 48S complex.

**Figure 2.**
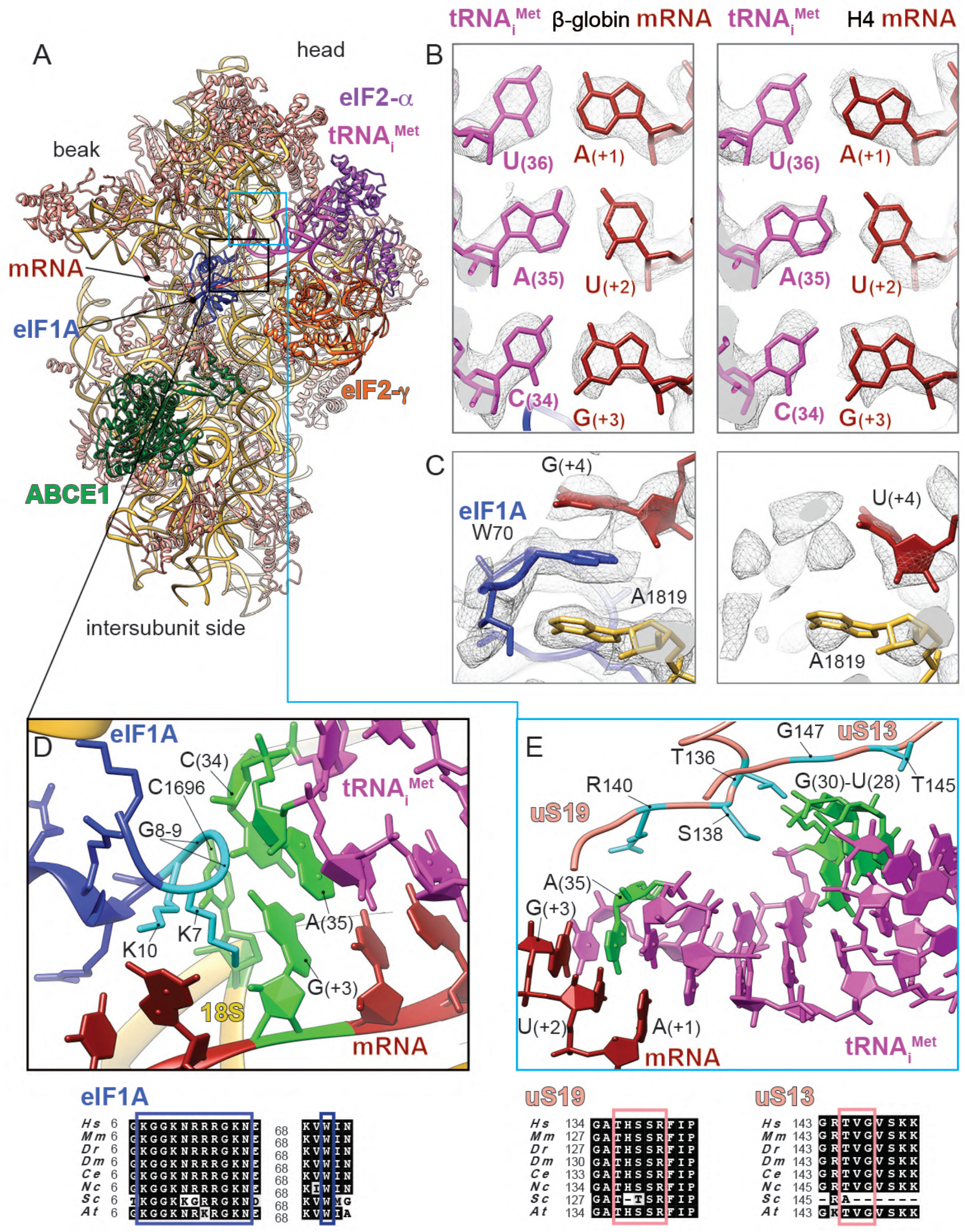
Key interactions surrounding the start-codon recognition sites in β-globin and H4 LS48S ICs. (A) Ribbon representation of the atomic model of β-globin LS48S IC viewed from the intersubunit side. (B) Codon:anticodon base-pairing view in both mRNA complexes; left: β-globin, right: H4. (C) eIF1A (in skyblue) interaction with the mRNA in the β-globin IC (left panel), compared to the corresponding region in the H4 IC, which is mostly free of eIF1A (right panel). (D) Close-up on the eIF1A N-terminal tail (coloured in cyan) showing its intricate interactions with tRNA and mRNA; stacking of C1696 on tip of tRNA ^Met^. The nucleotides involved in the interactions are coloured in green. (E) Interaction network of the tRNAi with ribosomal proteins uS13 and uS19 (coloured in salmon). Residues involved in the interactions are coloured in cyan in uS13 and uS19 and in green in the tRNAi. For eIF1A, uS13 and uS19, sequence alignments of the concerned interacting regions from eight representative eukaryotic species are shown below the panels in black boxes and the described residues are indicated by coloured frames (Hs : *Homo sapiens*, Mm : *Mus musculus*, Dr : *Danio rerio*, Dm : *Drosophila melanogaster*, Ce : *Ceanorhabditis elegans*, Nc : *Neurospora crassa*, Sc: *Saccharomyces cervisiae*, At: *Arabidopsis thaliana*).

C1696 of 18S rRNA is stacked on the C(34) base at the very tip of the tRNA_i_^Met^ ASL that is paired to G(+3) of both β-globin and H4 mRNA (Figure 2D, shown only for β-globin complex). This contact between C1696 and C(34) is also found in the yeast partial 48S pre-initiation complex (py48S IC) (Hussain et al., 2014) and it occurs even in the absence of any mRNA (des Georges et al., 2015). This stacking interaction may partly explain the difference in recruitment of initiator tRNA between bacteria and eukaryotes. In bacteria, the initiator tRNA is recruited directly at the P-site-accommodated start codon, whereas in eukaryotes, the tRNA_i_^Met^ is recruited at the pre-initiation stage of the complex before the attachment of mRNA into its channel. The tRNA_i_^Met^ ASL also interacts with the C-terminal tails of 40S ribosomal head protein uS19 (Figures 2E and S3B) through its Arg140 that contacts A(35).

We then compared the overall conformation of the 40S between both complexes and we observed that in the β-globin IC the head of the SSU is tilted downwards by ∼2° and swivelled toward the solvent side by ∼3° when compared to its counterpart in the H4 IC (Figure S2C). We attribute these subtle conformational changes to the dissociation of eIF1A in H4 LS48S IC, after the start-codon recognition, due to the loss of contacts between eIF1A and the 40S head.

### Interaction network of the Kozak sequence (−4 to +4) with 40S and initiation factors

The (+4) position, mainly occupied by a G in eukaryotic mRNAs, plays a pivotal role (Kozak, 1984, 1986). Our reconstructions demonstrate the structural importance of this position to both mRNAs. In the β-globin LS48S IC, the highly conserved Trp70 from eIF1A is trapped between the mRNA G(+4) position and the A1819 from h44 18S rRNA of the A-site by stacking interactions (Figure 2C). Interestingly, the interaction of the (+4) mRNA position with h44 was shown by cross-linking studies (Pisarev et al., 2006). Our β-globin LS48S IC structure also shows the proximity of uS19 C-terminal tail to (+4) mRNA position, which can also be corroborated by several cross-linking studies (Bulygin et al., 2005; Pisarev et al., 2008, 2006). In H4 mRNA a U is at position (+4), therefore the stacking interaction with eIF1A appears weaker than when a G is present. Moreover, nucleotides A1818 and A1819 have even more scant densities, indicating their undetermined conformations probably linked to this poor stacking (Figure 2C). Our reconstructions therefore suggest the structural importance of the (+4) position in the interaction with eIF1A.

Another crucial position in the Kozak consensus sequence is at (−3), often occupied by an adenine (Figure 1A). This nucleotide in both complexes shows several contacts with ribosomal proteins and initiation factors, including salt-bridge interaction between A(−3) base and a side chain of Arg55 from domain 1 (D1) of eIF2α (Figure 3A), which was reported previously in the py48S IC structure (Hussain et al., 2014; Llácer et al., 2018). However, in the yeast structure the A(−3) base is in the *syn* conformation and in both our mammalian ICs the adenine is in the *anti* conformation. Noteworthy, the near-cognate yeast mRNA present in the py48S IC structure (Llácer et al., 2018), contains adenines at the positions (−1) and (−2), which in principle could create an ideal stacking context for the A base in (−3), thus explaining this difference in conformation compared to our mRNAs where these positions are occupied by two cytosines. The (−3) position further interacts with the G957 nucleotide at the 40S platform (Figure 3A), highlighted in earlier studies (Demeshkina et al., 2000). In addition, cross-linking studies of reconstituted mammalian PIC previously demonstrated that eIF2α and uS7 interact with the (−3) nucleotide, and uS7 with the (−4) nucleotide (Pisarev et al., 2008, 2006). The interaction of uS7 through its β-hairpin was also suggested in the py48S IC structure, due to their proximity in space (Hussain et al., 2014; Llácer et al., 2018). However, in our structures, this interaction cannot be confirmed since the electron density at this specific region is very disperse, probably because of the flexibility of this part of uS7 (see Discussion, Figure S5A) compared to its other parts.

**Figure 3.**
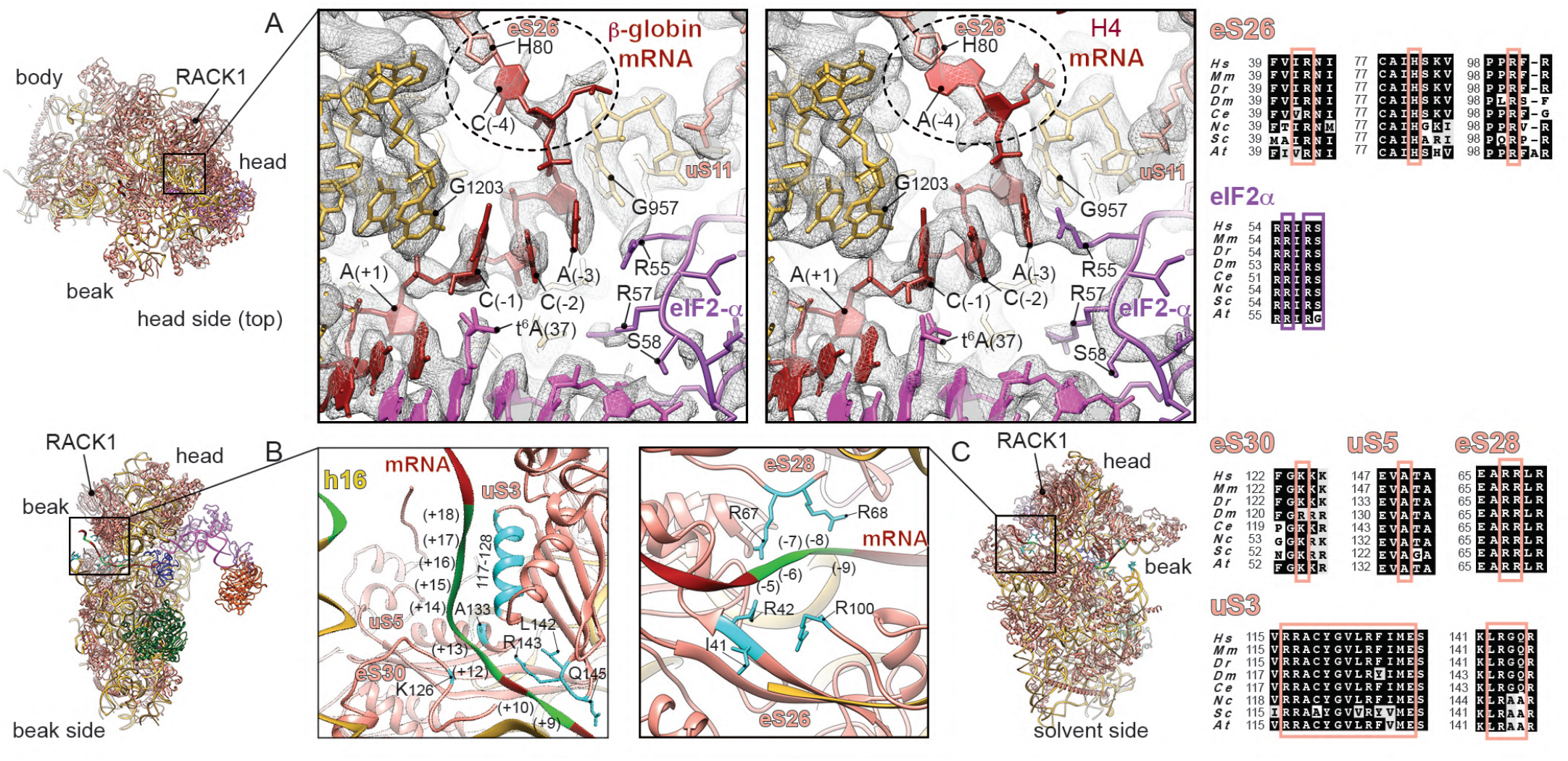
Kozak and beyond Kozak interaction networks in β-globin and H4 LS48S ICs. (A) Close-up of the interactions of upstream start-codon nucleotides top-viewed from the head side in the β-globin (left panel) and H4 (right panel) ICs with ribosomal proteins eS26 and uS11, as well as eIF2α D1 domain, tRNAi and 18S rRNA. mRNA (−4) position contact with His80 of eS26 is highlighted in dashed line circle. The distances between atom N1 of His80 eS26 and amine groups of A(−4) (β-globin) and C(−4) (H4) are 3.7 Å and 3.2 Å, respectively. G1203 and G957 of 18S rRNA stacking and interaction with C(−1) and A(−3), respectively, of both mRNAs are shown. (B) mRNA entry channel seen from the beak side with close-up on the interactions with uS3, eS30 and h16 of 18S rRNA. (C) mRNA exit channel seen from the solvent side with close-up on the mRNA contacts with ribosomal proteins eS26 and eS28. (B) and (C) panels are shown on an example of β-globin LS48S IC. The nucleotides involved in the interactions are indicated in green and residues in cyan. Respective sequence alignments are shown in black boxes from eight representative eukaryotic species on the right of the figure panels.

Position (−4) of both mammalian mRNAs interacts with ribosomal protein eS26 through its His80. However, we have found that in the case of the β-globin mRNA, position (−4) is a cytosine and appears to interact mildly with eS26 His80 (Figure 3A, left panel), as its weak density suggests. Whereas when this position is an adenine, like in the H4 mRNA, a stronger stacking interaction occurs, which could further participate in stabilizing the mRNA in its channel (Figure 3A, right panel). Consequently, the mRNA in this latter case adopts a slightly different conformation. A possible result of this difference is the observed tighter interaction with eIF2α Arg55 residue from domain D1 (Figure 3A), as its density is better defined in H4 than in β-globin.

Finally, upstream residues near the start codon in our complexes are in contact with 18S rRNA including G1203 from the head rRNA which interacts with the phosphate of A(+1) and stacks with C(−1) of both mRNAs (Figure 3A).

### eIF1A interaction with the β-globin mRNA sequence and the 18S rRNA

In addition to the above-mentioned contact with the start codon and G(+4), eIF1A can potentially establish several interactions at more distal positions in the β-globin mRNA sequence, closer to the mRNA entrance channel. Indeed, Arg12, Lys67 and Lys68 in eIF1A are in close proximity to C(+7), G(+6) and U(+5) (Figure 4A). eIF1A NTT also interacts with the 18S rRNA (Gly9 and Lys10 with C1696; and Lys16 and Asn17 with C1327) (Figures 2D and 4A). Other contacts involve the loops of the eIF1A OB domain with the 40S near the A-site (Figures 4B-4D): namely, Asn44 and Arg46 are in contact with A1817-A1819 and C1705 from h44 of 18S rRNA (Figure 4C); moreover Lys64 and Arg62 contact G604 and C605 of h18 18S rRNA (Figure 4D). In addition, Arg82, Tyr84 and Gln85 of eIF1A contact Glu58, Leu91 and Gly56 of ribosomal protein uS12 (Figure 4D); finally, Asp83 is in contact with Arg82 of eS30 (Figure 4D). Putting together, the above-mentioned interactions might depend on the mRNA sequence and perhaps they can have influence on the stability the cognate start-codon duplex with its anticodon by the NTT of eIF1A (residues Lys7, Gly8, Gly9 and K10).

**Figure 4.**
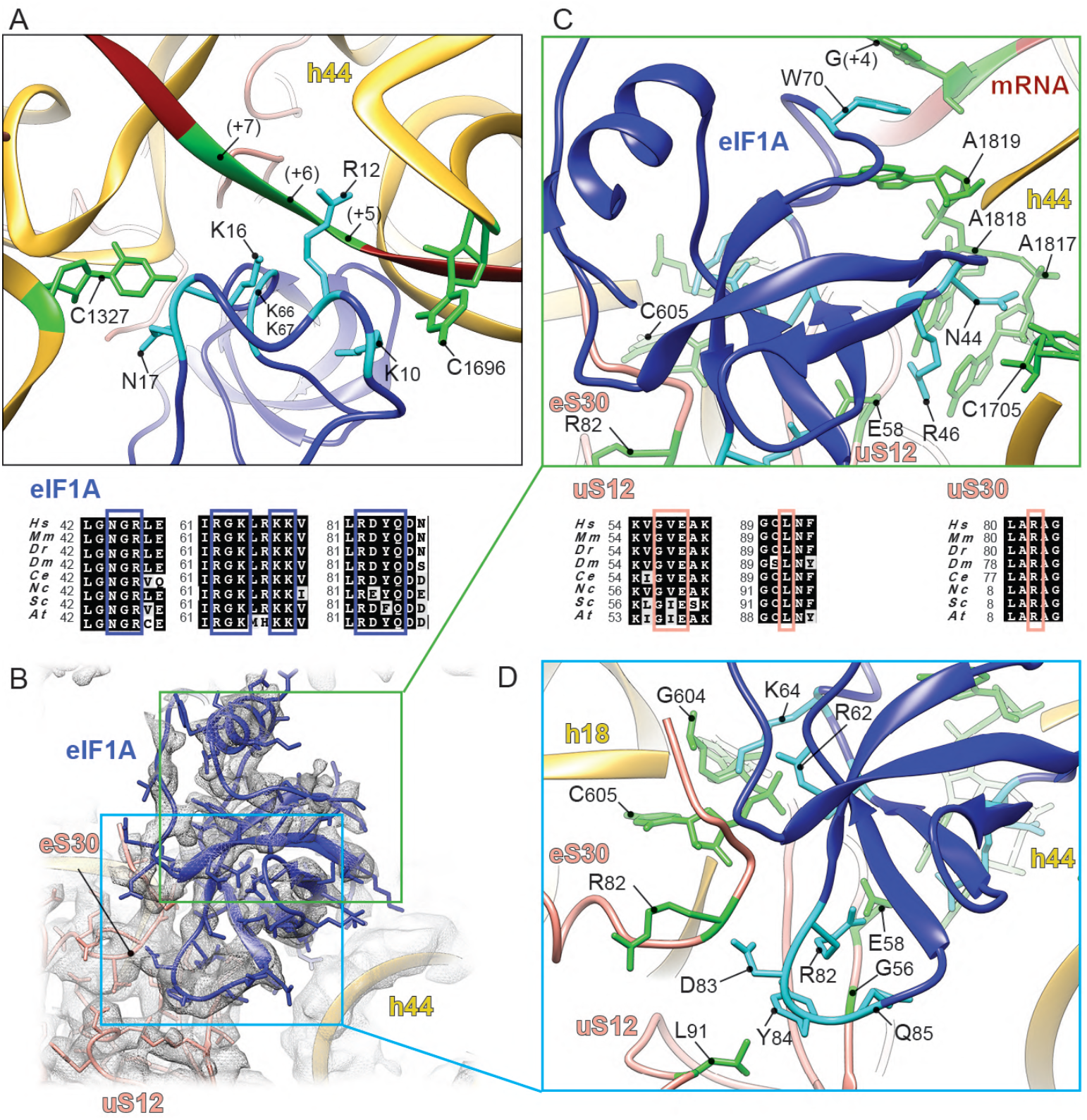
eIF1A interactions in the A-site of β-globin LS48S IC. (A) eIF1A (in dark blue) N-terminal tail interactions with mRNA of downstream start-codon nucleotides and tRNAi. The nucleotides involved in the interactions are indicated in green and residues in cyan. (B) eIF1A OB-domain interactions with mRNA and 40S. (C) Close-up on interactions of eIF1A (in dark blue) with h44 of 18S rRNA (nucleotides are coloured in green). (D) Zoom in on eIF1A (in dark blue) interactions with h18 of 18S rRNA (in gold) and ribosomal proteins uS12 and eS30 (in salmon). The nucleotides and residues of uS12 and eS30 involved in the interactions are indicated in green and eIF1A residues in cyan. Respective sequence alignments are shown in black boxes.

Noteworthy, eIF1A NTT was shown to interact with eIF5 (Luna et al., 2012, 2013), but because of its clear involvement in the start codon:anticodon duplex, we suggest that this eIF5 interaction occurs during the pre-initiation phase and very shortly after the recruitment of the mRNA.

### mRNA interactions with the 48S beyond the Kozak sequence

The mRNA density at distal positions from the Kozak sequence appears disperse when filtered to high-resolution, suggesting an overall flexibility at both the entrance and the exit of the channel (local resolution of ∼6 to ∼9 Å). Nevertheless, several contacts can be observed at the entrance and exit sites of the mRNA channel of the β-globin and H4 LS48S ICs. These interactions are common to both complexes and could be more site-specific than they are sequence-specific.

At the entrance of the mRNA channel during this late stage of the initiation process, the mRNA extensively interacts with conserved residues of the 40S ribosomal proteins uS3 and eS30, and with rRNA h16 in positions spaning from +10 to ∼ +20 through ionic and hydrophobic interactions (Figure 3B). For instance, the conserved Arg117 of the head protein uS3 contacts the mRNA at the channel entrance. This residue was recently indicated as important for stabilizing the P_IN_ closed state of the 48S in yeast IC (Llácer et al., 2018) and for the initiation accuracy in the presence of suboptimal Kozak sequence by *in vivo* assays in yeast (Dong et al., 2017). The contribution of this charged residue of uS3 contacting the mRNA is partially corroborated by cross-links in a previous study (Pisarev et al., 2008). More globally, charged amino acid residues from uS3 helix α (residues 117-128) are in close proximity to nucleotides from positions (+14) to (+18) (Figure 3B). Moreover, residues from a β-hairpin (residues 142-146) can potentially contact bases of the nucleotides C(+9) and C(+10) of the mRNA, forming hydrophobic and salt-bridge interactions. For ribosomal protein eS30, Lys126 is in close distance to the bases of G(+12) and A(+13). The proximity of A(+13) of mRNA to Ala133 of uS5 can also be noted (Figure 3B).

On the other side at the mRNA exit channel, we can observe the exit of both β-globin and H4 mRNAs from their respective 48S ICs (Figure 3C). The 5’ untranslated region (5’UTR) for β-globin mRNA is substantially longer than for H4 (50 nt and 9 nt, respectively) (Figure 1A, see Methods). We compared the mRNA exit channels of both complexes below the ribosomal head protein RACK1, which unambiguously shows the expected larger 5’ UTR for β-globin LS48S IC compared to H4 (Figure S2B). We were able to spot several possible contacts of ribosomal proteins at the exit site with mRNA nucleotides in both LS48S IC, including eS28 (Arg66 with A(−5), and Arg67 with A(−7)) as well as eS26 (Ile41 with C(−8), Arg42 with A(−9) and Arg100 with both these nucleotides) (Figure 3C), in agreement with previous cross-linking results (Pisarev et al., 2008, 2006).

### Interactions of 48S initiation complex with the tRNA_i_^Met^

The overall accommodation of the mammalian ASL resembles its yeast counterpart found in the P_IN_ state (Hussain et al., 2014; Llácer et al., 2018). The 48S IC-tRNA_i_^Met^ interaction network is summarized in the Figure S3B. In both ICs, we can observe a density attached to A(37), in which we can model the threonylcarbamyol group forming a t^6^A modification (Figures 3A and 5A). This modification mediates the binding of t^6^A(37) to the 2’OH of C(−1) in the mRNA, and therefore can further stabilize the start-codon recognition. It is tempting to suggest that C(−1):t^6^A(37) interaction is required for efficient translation in mammals. This mRNA C(−1) position is conserved in higher eukaryotes, as revealed by quantitative sequence analysis (Grzegorski et al., 2014), and forms part of the Kozak sequence. Interestingly, the electron density for this modification is even stronger in the H4 mRNA complex than in β-globin, even at a lower resolution (3.5Å). We therefore suspect that this interaction could be more important in the case of suboptimal Kozak sequences, where this modification could compensate for the loss of some interactions with the 48S IC, compared to a stronger Kozak mRNA. The same interaction in the case of A at (−1) position is not excluded, however its nature and conformation will be different.

**Figure 5.**
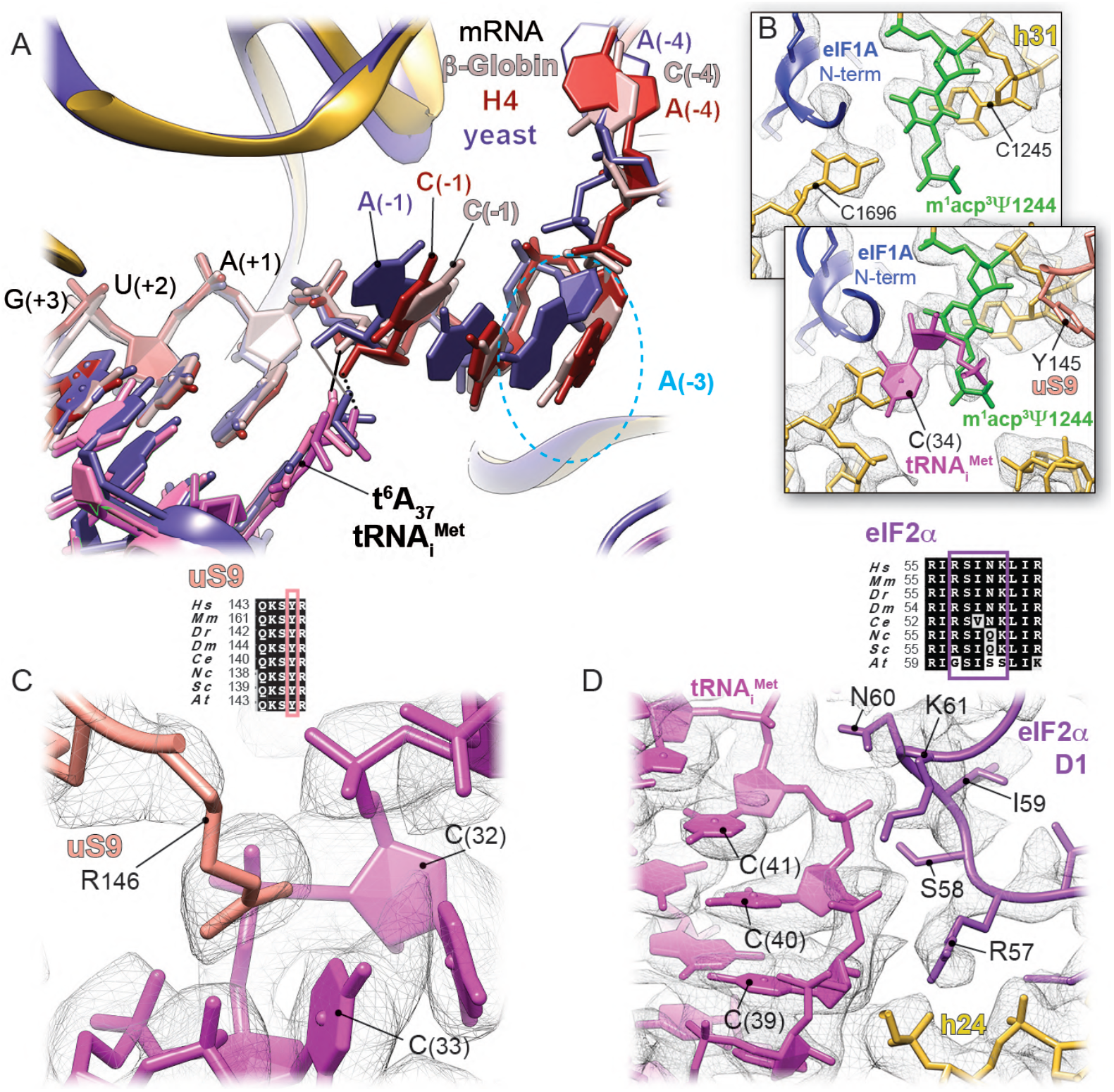
Initiator tRNA anticodon stem-loop (ASL) interactions with the LS48S IC. (A) Comparison of the tRNAi modified t^6^A(37) interaction with mRNA (−1) position in mammalian β-globin (in light pink) and H4 (in red) and yeast (Llácer et al., 2018) (dark purple) initiation complexes. The contact with (−1) mRNA position is labelled by black solid line, black dots and grey solid line for β-globin, H4 and optimal yeast mRNA sequence, respectively. In dashed skyblue circle, comparison of the conformation of highly conserved A(−3) mRNA position: *anti* in mammalian IC and *syn* in yeast. (B) Interaction of the modified m^1^acp^3^Ψ1244 of the 18S rRNA (coloured in green, overlapping bottom panel) with the C(34) of the tRNAi and ribosomal protein uS9 (overlapping top panel). (C) Close-up on the interactions of C(32) and C(33) with Arg146 of uS9. (D) Close-up on the interaction of the ASL cytosines from the conserved G-C base-pairs with eIF2α domain D1. Respective sequence alignments are shown in black boxes and interacting residues in coloured frames.

Despite the universal presence of the t^6^A hypermodified base in all organisms, and a crucial role in translation efficiency (Pollo-Oliveira and De Crécy-Lagard, 2019; Thiaville et al., 2016), it has only recently been shown that the modification directly contributes to the AUG recognition accuracy. In the py48S-eIF5 IC structure at a resolution of 3.5Å (Llácer et al., 2018), t^6^A of the initiator tRNA^Met^ was suggested to enhance codon:anticodon base-pairing by interacting with A(−1) and by a stacking on the downstream base-pair involving A(+1). Thanks to our mammalian LS48S IC at 3.0Å, we can clearly observe the threonylcarbamyol group in a different conformation, placing the carboxyl group within hydrogen-bonding distance (2.7Å) from the 2’OH group of C(−1) (Figure 5A). However, this modification does not appear to stack over the downstream base-pair as previously suggested. Noteworthy, in yeast there is a preference for an A at position (−1) (Dvir et al., 2013; Kozak, 1986), while it is a C(−1) in mammals.

Furthermore, C(34), that is a part of the anticodon, is stabilized by the modified U1244 (U1248 in human) of helix31 of rRNA (m^1^acp^3^Ψ, 1-methyl-3-(3-amino-3-carboxypropyl) pseudouridine) (Maden, 1990; Taoka et al., 2018) (Figure 5B), which was previously reported in the py48S-eIF5 (modified U1119) (Llácer et al., 2018). In addition, the neighbouring nucleotides, C(32) and C(33), are in contact with C-terminal arginine Arg146 of ribosomal head protein uS9 (Figure 5C).

Aside from its role in the codon:anticodon stabilization, uS19 together with uS13 were found to contact other parts of the tRNA_i_^Met^ through their highly conserved C-terminal tails (Figures 2E and S3B). Thr136 side chain of uS19 interacts with the guanine backbone of three conserved G–C base pairs in ASL that are crucial for stabilization of the initiation complex in eukaryotes (Dong et al., 2008). Residues Thr145 and Gly147 of uS13 are in contact with the phosphate groups of U28 and G29 (Figure 2E). Consistent with previous reports (Hussain et al., 2014; Llácer et al., 2018), several residues from domain D1 of eIF2α (Arg57, Ser58, Asn60 and Lys61) interact with the cytosines backbones of three conserved G–C base pairs of the ASL (phosphate groups of C39-41) (Figures 5D and S3B).

### ABCE1 binding to the initiation complex is NTP-dependent

ABCE1 (named Rli1 in yeast) is a conserved NTP-binding cassette ABC-type multi-domain protein that plays a role in translation initiation as well as translation termination and ribosome recycling (Becker et al., 2012; Heuer et al., 2014; Khoshnevis et al., 2010; Pisarev et al., 2010; Shoemaker and Green, 2011; Young et al., 2015). It contains two nucleotide-binding domains (NBDs), where the two NTP molecules bind. Its N-terminal NBD contains two iron-sulphur clusters [4Fe-4S] 2+ (Barthelme et al., 2007, 2011; Karcher et al., 2008). In our β-globin LS48S IC structure, the NTP-binding cassette of ABCE1 displays lower local resolution (between 3.5 and 5Å, Figures 6A and S1D) compared to the average resolution, likely due to the flexibility of the NBDs. In H4 LS48S IC structure we observe only a residual density of ABCE1 (Figure S1L), which can be caused by a slightly different conformation of h44 18S rRNA in the absence of eIF1A.

**Figure 6.**
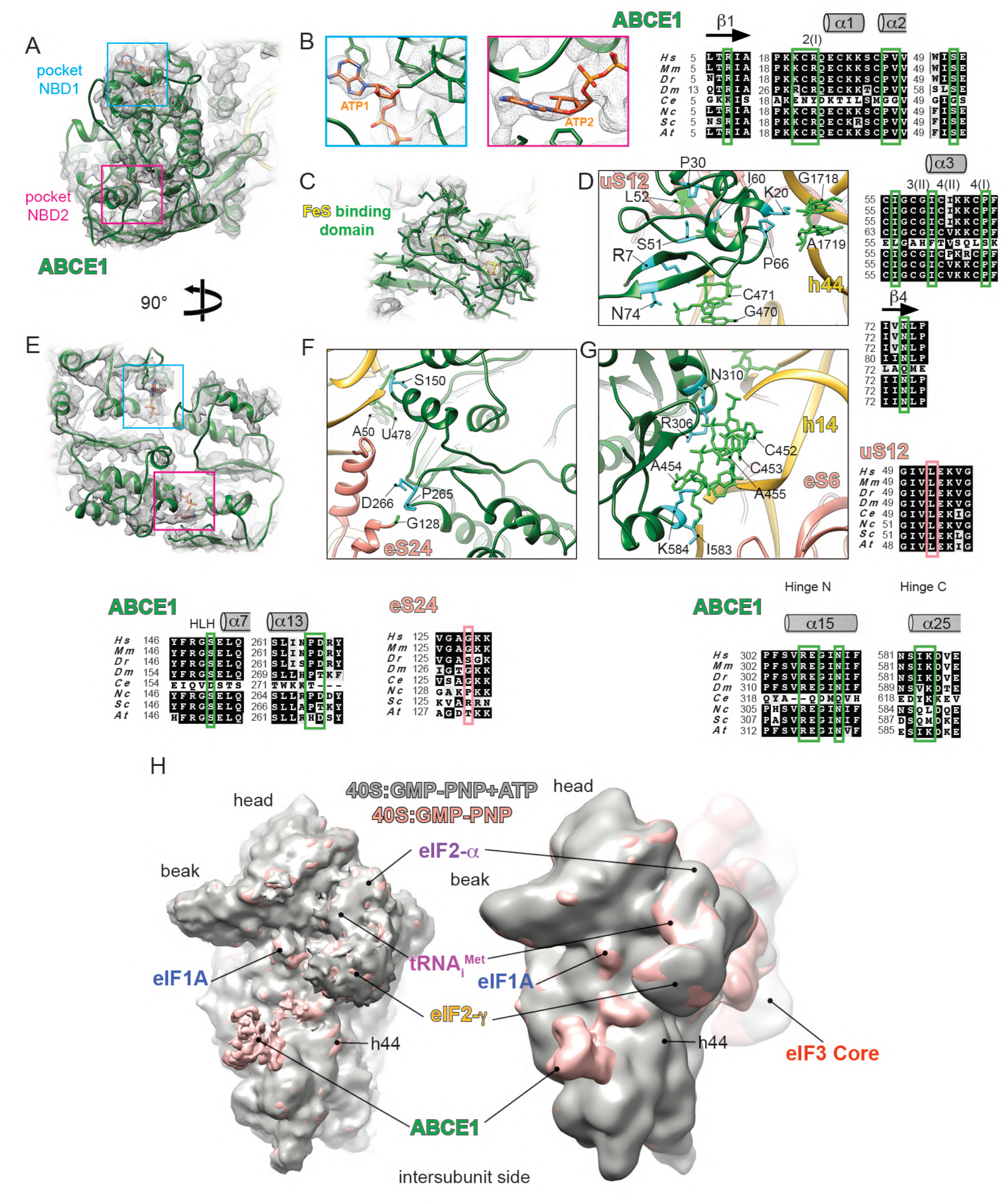
Atomic model of ABCE1 in the LS48S IC. (A) and (E) Ribbon representations of ABCE1 (in green) in its electron density with NTP-binding pockets framed in blue (NBD1) and pink (NBD2) seen from different side views. (B) Blowups on the NTP pockets, NBD1 (left blue) and NBD2 (right pink). Although in the purification conditions GMP-PNP was used, ATP molecules were modelled in the electron densities obtained. (C) Mixed ribbon and stick representation of Fe-S binding domain atomic model fitted into its electron density. (D) Close-up of Fe-S binding domain interactions with 40S ribosomal protein uS12 and h44 of 18S rRNA. (F) Close-up on NBD1 interactions with the 40S. (G) Close-up on NBD2 interactions with the 40S. The nucleotides involved in the interactions are indicated in light green and protein residues in cyan. Respective sequence alignments are shown in black boxes. (H) Comparison between the β-globin (pink surface) and β-globin•GMP-PNP+ATP (grey surface) LS48S ICs reconstructions without (left superimposition) and with (right superimposition) eIF3. The panel shows that the addition of ATP triggers the dissociation of ABCE1, probably after its hydrolysis. No further conformational changes between both complexes, with and without ATP, can be detected.

The iron-sulphur (Fe-S) binding domain presents a higher local resolution than the NBDs (between 3 and 4Å, Figures 6B and S1D). Therefore, we can clearly distinguish the Fe-S clusters in the cryo-EM density map as well as the presence of two bound nucleotides that probably represent GMP-PNP that was used to stall the initiation complexes by blocking eIF2γ (Figures 6A-6C).

NBD1 contacts five nucleotides in the 18S rRNA helix14 (U478, A455, A454, C453 and C452) through residues located in a helix-loop-helix motif (Ser150) and in the hinge-N (Arg306 and Asn310) (Figures 6F-6G and S4A). NBD2 also makes contacts to nucleotides A455, A454, C453 through the hinge-C residues Lys584 and Ile583, conserved in higher eukaryotes (Figures 6G and S4A). Residues Arg566 and Arg567 from NBD2 are in close proximity to the 18S rRNA and can potentially be involved in the interaction with the latter, as suggested by their conservation (Figure S4A). Moreover, NBD1 residues Pro265 and Asp266 interact with residues from the C-terminal helix of ribosomal protein eS24 such as Gly128 (Figure 6F).

As for the Fe-S binding domain, it interacts with the 18S rRNA (Arg7 from strand β_1_ to C471, Lys20 to G1718, Pro66 to A1719 and Asn74 to G470) (Figure 6D). In addition, ABCE1 interacts with ribosomal protein uS12 residues Ile50, Leu52 and Ile75 via a hydrophobic pocket formed by helices α_2,_ α_3_ and cluster (II) (residues Pro30, Val31, Ile56, Ile60) (Figure 6D). This is consistent with previous structural and crosslinking studies in yeast and archeal complexes (Heuer et al., 2017; Kiosze-Becker et al., 2016; Nürenberg-Goloub et al., 2020).

In order to investigate the effect of ABCE1 binding on the structure and composition of the LS48S IC, we purified β-globin complexes using our choice strategy (Simonetti et al. 2016 and this work) supplemented with 10 mM of ATP, thus taking advantage of the ability of ABCE1 to hydrolyse ATP, in contrast to the obligate GTPase eIF2. The β-globin-ATP 48S IC was then analysed by cryo-EM and yielded two main LS48S IC reconstructions at ∼14Å and ∼10Å, that differ in the presence and absence of eIF3, respectively (Figures 6H and S1X,Y). The addition of ATP most likely causes the replacement of the GMP-PNP molecules in the NBD pockets by ATP molecules that was then hydrolysed by ABCE1. Our structures clearly reveal the dissociation of ABCE1 as a result of ATP addition, consistent with recent structural and biophysical studies (Gouridis et al., 2019; Heuer et al., 2017; Kiosze-Becker et al., 2016). Aside of the absence of ABCE1, the global structure of the β-globin-ATP LS48S IC is identical to its higher-resolution counterpart without ATP (Figure 6H), thus very likely excluding a direct active role of ABCE1 in the assembly of the initiation complex. It is reasonable to assume that in the cell ABCE1 undergoes on/off cycles to the IC in an ATP-dependent manner, as we have previously suggested (Mancera-Martínez et al., 2017). However, these results do not contradict the demonstrated function for ABCE1 as an anti-ribosomal-subunit association factor.

### eIF3 in the late-stage 48S initiation complex

Nearly 15% (∼5% of the total particles count) of the particles corresponding to the β-globin LS48S IC structures contain a density for eIF3 at the solvent side. After extensive particle sorting and refinement, a reconstruction of the β-globin LS48S IC showing eIF3 was obtained at a resolution of ∼3.6Å (Figures S1I–S1O), thus allowing verification of the recognition of the start codon for this specific class (Figures 7A–7B). eIF3 a and c subunits, i.e. those that bind directly to the 40S, are mostly resolved at a resolution ranging from 3.5 to 4.5Å, enabling a model of the exact residues in interaction with the 40S subunit (Figures 7C–7G). We find that eIF3a in β-globin LS48S IC shows several contacts with 40S ribosomal body proteins, with eIF3a residues Asn10, Lys13, Arg14 and Phe18 interacting with eS1 residues Asp77, Asn76, Asp191 and Pro190, respectively (Figures 7F–7G). In addition, eIF3c residues Asn388, Arg340, Asn384, Gly341, Lys343 and Arg450 contact the 40S ribosomal protein eS27 via residues Glu75, Thr61, Gln65 and Cys59 (Figure 7E). Moreover, residues Lys342, Lys343, Thr391 and Tyr392 interact with nucleotides G925, C1112 and U1116, two latter being a part of the apical loop of expansion segment 7 (ES^7^).

**Figure 7.**
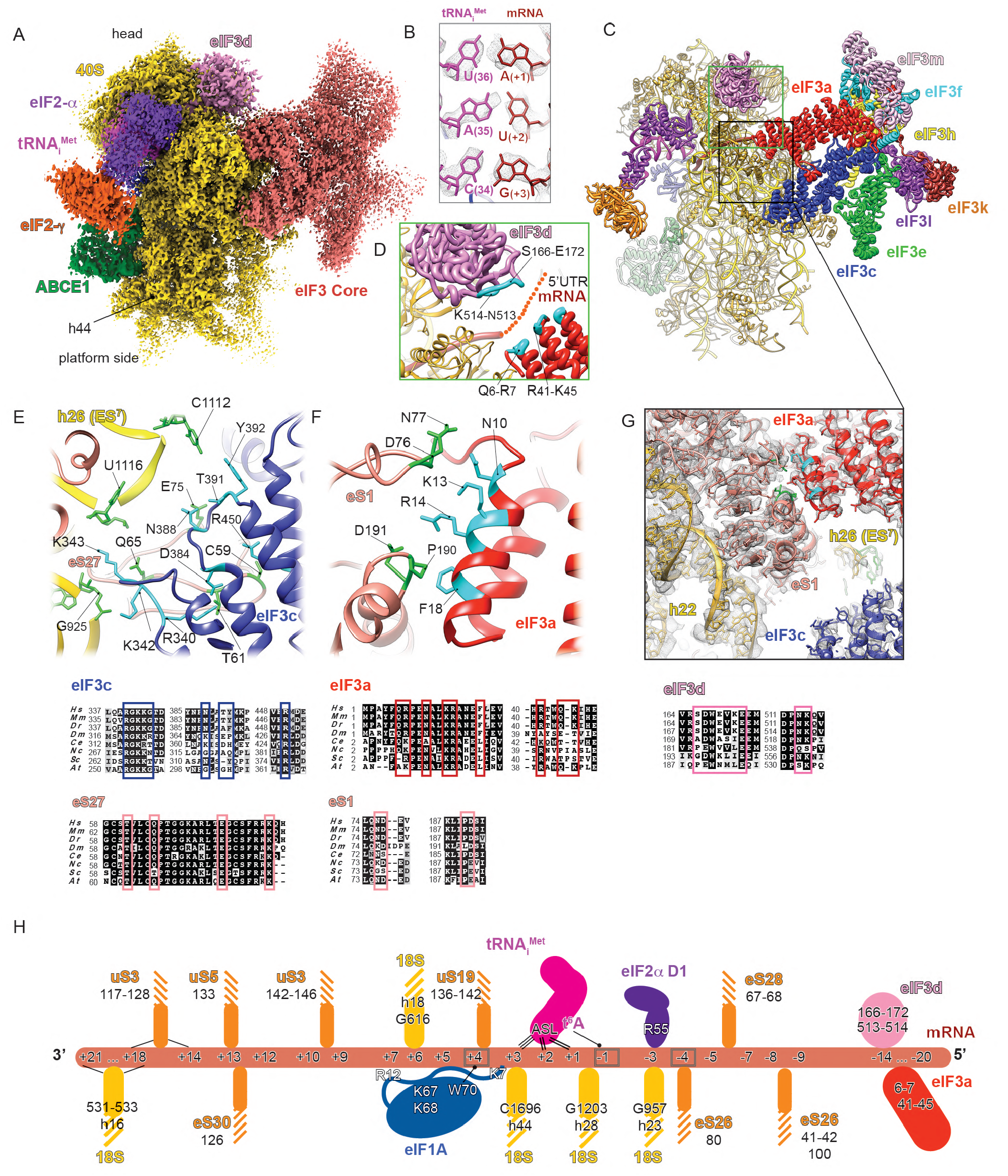
Cryo-EM reconstruction of the eIF3-containing class of the β-globin LS48S IC. (A) Segmented map showing electron density of the eIF3 core (in rose) attached to the 48S viewed from the platform side. (B) Codon:anticodon base-pairing view in LS48S-eIF3 ICs (identical to both β-globin and H4). (C) Ribbon representation of the atomic model of LS48S-eIF3 IC seen from the platform side. (D) Blowup on the mRNA channel exit, seen from the platform side. mRNA 5’UTR cannot be modelled because of the low local-resolution of the cryo-EM reconstruction in this region, therefore we propose an extrapolation of the mRNA 5’UTR trajectory showing possible interactions with eIF3 d and a subunits (residues coloured in cyan). (E) Close-up on the eIF3c (in navy blue) interaction with 40S: h26 (ES^7^) of 18S rRNA and eS27. (F) Close-up on the eIF3a (in coral) interaction with eS1. (G) Mixed ribbons and sticks representation of eIF3 core interactions with 40S. The nucleotides involved in the interactions are indicated in chartreuse and protein residues in cyan. Respective sequence alignments are shown in black boxes with interacting residues highlighted by coloured frames. (H) Summary of the mRNA interactions with mammalian LS48S IC. The ribosomal proteins are coloured in orange and 18S rRNA elements in yellow. The mRNA contacts critical for recognition of optimal or suboptimal Kozak context are highlighted by grey frames.

The eIF3d subunit structure is at lower resolution as compared to the eIF3 octamer core. Nevertheless, secondary structure elements can clearly be depicted when filtered to a lower resolution (6Å), which enabled the fitting of its partial crystal structure in our density (Lee et al., 2016). The modelled eIF3d displays contacts with several ribosomal head proteins: helix α_12_ contacts the N-terminal loop of uS7; the loop between β_9_ and β_10_ contacts RACK1 (loop located between strands β_6D_ and β_7A_); and β-strand loops and the “RNA gate” insertion contacts eS28 (strand β_3_, loops between helices α_8_ and α_9_).

In spite of the low local-resolution of this particular subunit, our structure provides clues to the demonstrated interactions of eIF3d and eIF3a with the mRNA 5’UTR (Figure 7D). Thus, shown by the residual electron density traces, the 5’UTR of numerous mRNAs such as β-globin can possibly interact with different parts of eIF3d: N-terminal loop and strand β_2_ (residues S166 to E172) and a loop between β_11_ and β_12_ (residues Asn513 and Lys514) (Figure 7D). These residues of eIF3d are better conserved in higher eukaryotes, indicating a possible species-specific regulation. An interaction with 5’ RNA terminus recognition motif was also previously reported (Lee et al., 2016). More insight into the interaction patterns of eIF3d with different mRNA 5’UTRs in the context of translation initiation will be an important goal in future studies.

Finally, the mRNA 5’UTR of β-globin can also interact with eIF3a (residues Gln6, Arg7, Arg41, Gln44 and Lys45) (Figure 7D). Similarly to eIF3d, these residues are mainly conserved among higher eukaryotes (Figure 7G). It was previously shown that the eIF3a–PCI domain (a domain with a common fold for proteasome, COP9, initiation factor 3) is critical for stabilizing mRNA binding at the exit channel (Aitken et al., 2016). However, because of the low local resolution of the β-globin mRNA 5’UTR in our reconstruction, we do not exclude other patterns of interaction. H4 LS48S IC also shows the residual presence of eIF3 core, however particle sorting reveals a reconstruction containing eIF3 (Figure S1W) at only intermediate resolution because of the low number of particles.

## DISCUSSION

Our cryo-EM structures reveal in detail the accommodation of two native mRNA sequences encoding either β-globin or histone 4 in the context of the late-stage mammalian IC. In the presented IC structures, we did not identify any density corresponding to eIF1. Combined with codon:anticodon complex formation and several conformational changes characteristic of the stage after the start-codon recognition, we have dubbed our complexes “late-stage 48S IC (LS48 IC)”. Initiation on β-globin and H4 mRNA may undergo different regulatory processes as previously reported (Martin et al., 2011, 2016), however in our structures we only analyse the mRNA nucleotide interactions of the Kozak sequences (Figure 7H and Table S2) without dwelling on the exact regulation mechanism that may be in part influenced by the different interaction patterns that we observe. We therefore believe that our two archetype mRNA sequences are representative of native cellular mRNAs incorporating different Kozak sequences, as their observed interactions are purely the result of sequence differences, and unrelated to specific regulatory pathways.

In the Kozak sequence, the position (+4) appears to play a role in both mRNAs, where it is a G in β-globin and U in H4. At this position, the crucial interaction with Trp70 of eIF1A appears to be weaker in the case of H4 IC when compared to β-globin IC, as indicated by our MS/MS normalized spectral counts and the cryo-EM reconstructions. We suggest that the poor abundance of eIF1A in the H4 LS48S IC cryo-EM reconstruction (Figures 1, S1L and S2A) might not suggest a negligible role for this initiation factor in suboptimal Kozak consensus mRNAs during scanning. Rather, it simply shows its weaker interaction in the complex after start-codon recognition, supported by our semi-quantitative MS/MS analysis. As expected, the biggest decrease in spectral counts was observed for eIF1A in the H4 LS48S IC compared to its β-globin counterpart (Figure 1B). The weak above-mentioned stacking interaction with mRNA purine at (+4) position is likely to be the reason of the affinity drop and subsequent weaker interaction between eIF1A and the 48S in the case of H4, after scanning and the recognition of the start codon. This observation is consistent with the suggested special translation initiation mechanism for H4 (Martin et al., 2011, 2016). It was suggested that H4 mRNA undergoes an unconventional “tethering mechanism”, where the ribosomes are tethered directly on the start codon without scanning (Martin et al., 2011). This particular mechanism was proposed to occur thanks to the presence of two secondary structure elements present downstream of the start codon in H4 mRNA (position +19), contacting h16 of rRNA (Martin et al., 2016). In this region, our H4 LS48S IC structure shows disperse densities that cannot be interpreted. We believe that eIF1A is present in the H4 complex during pre-initiation and the short scanning process, and only after the recognition of the start codon the affinity can be affected by the mRNA Kozak context.

Regarding the N-terminal tail of eIF1A, it is present in the A-site starting from the scanning process, as shown by our structure and also yeast 43S PIC and closed-48S ICs (Llacer et al., 2015; Llácer et al., 2018). It is important to emphasize the binding of eIF1A on the H4 IC at a certain stage, because we can observe residual electron density for this initiation factor in the H4 LS48S IC (Figures S1L and S2A). This observation tends to validate the unconventional very short scanning mechanism proposed for H4 mRNA (Martin et al., 2011), yielding in the faster accommodation of the start codon as compared to β-globin mRNA. Nevertheless, in the early initiation steps eIF1A may interact with eIF5, as demonstrated by biophysical studies of *in vitro* purified proteins (Luna et al., 2013).

The importance of positions (+4) and (−3) of mRNA has been pointed out in previous studies (reviewed in (Kozak, 1989)), however the significant involvement of position (−4) in mammals was not highlighted. Similarly to py48S-eIF5 IC (Llácer et al., 2018), our structures show that when this position is A, it can be stabilized by residue His80 of eS26. This residue is highly conserved in eukaryotes (Figure 3A), therefore showing a universal mode of interaction. The role of eS26 (and eS28) in the accommodation of the 5’ UTR was also highlighted by chemical cross-linking studies performed with 80S ribosome assembled on H4 mRNA (Martin et al., 2016). Remarkably, in the case of the yeast IC, it was shown that the interaction between the A nucleotide at the position (−4) and His80 of eS26 does occur through stacking (Llácer et al., 2018), in contrast to our study where we can show C and mainly A (−4) stacked below the His80 of eS26. This variation is probably due to the different sequence between our mRNA and those of yeast, where positions −1 to −3 are occupied by A, that favours more stacking between the nucleotides bases and consequently more twist (Figure 5A). Indeed, the kink in E/P site in py48S-eIF5 IC (Llácer et al., 2018) is sharper than in the case of mammalian 48S IC (Figure S3A). Interestingly, a previous cross-linking study in the context of the human 80S ribosome highlighted the eS26 binding to G and U nucleotides in the region (−4) to (−9) of the mRNA (Graifer et al., 2004). In a more recent biochemical study in yeast, it was shown that the mRNAs bound to the ribosomes depleted in eS26 (RpS26 in Saccharomyces) translate poorly when compared to those enriched in eS26 (Ferretti et al., 2017). It was also reported in the same study that RpS26 is necessary for preferential translation of mRNAs with A at (−4) position and not G, showing that the interaction is very specific and not simply purine/pyrimidine dependent. Therefore, the His80 eS26 recognition is likely optimal for mRNA sequences containing A at this position. In yeast, however, the nucleotide context surrounding the start codon is less critical but shares with mammals the importance of the (−3) position (Cavener and Ray, 1991; Kozak, 1986). Further studies on translation of mRNAs containing mutations at these positions will help unveiling the mechanism of scanning in mammals and will shed light on the leaky scanning mechanism.

The tails of uS13 and uS19 have been previously shown to make direct interactions with the ASL of the peptidyl-site-exit-site (P/E) tRNA in presence of elongation factor G-ribosome complex in a pre-translocation state in prokaryotes (Zhou et al., 2013). To the best of our knowledge, these proteins have not yet been reported to be particularly involved in the initiation process in eukaryotes. Nevertheless, our structures are supported by earlier crosslinking studies of human 80S ribosome showing that the tail of uS19 is located closer to the decoding site than that of prokaryotic S19 (Graifer et al., 2004). In the case of *S. cerevisiae* uS13 and uS19, the C-terminal parts are not conserved, compared to human protein homologues (Figure 2E) and they have never been observed to interact with the tRNA_i_^Met^ (Hussain et al., 2014; Llácer et al., 2018). Other fungi, such as *N. crassa,* possess very similar sequences to mammalian counterparts and probably would demonstrate similar interactions to tRNA_i_^Met^ as shown by our structures. Moreover, and in contrast to yeast, *N. crassa* possesses a mammalian-like eIF3. This is in line with the recent genome-wide mapping of Kozak impact in the fungal kingdom, showing the particularity of start-codon sequence context in *S. cerevisiae* compared to other fungi (Wallace et al., 2019).

The ribosomal protein uS7 is located close in space to the position (−3) of mRNA, but we cannot confirm its interaction with this nucleotide due to the lack of the density for this part of the protein. Noteworthy, this is the only flexible region in this protein structure (Figure S5A), containing the crucial β-hairpin in the case of bacterial and yeast initiation (Visweswaraiah and Hinnebusch, 2017; Wimberly et al., 1997). This region in yeast contains the glycine-stretch G<u>GG</u>G (residues 150-153), whereas in human it is G<u>RA</u>G (residues 129-132). Genetic experiments on single-point mutants of this β-hairpin demonstrated almost unchanged phenotype for human-like G151R and G152A mutations (Visweswaraiah et al., 2015), but the G151S mutation was lethal. More recent work showed by using genetic and biochemical approaches that uS7 modulates start-codon recognition by interacting with eIF2α domains in yeast (Visweswaraiah and Hinnebusch, 2017). The residues implicated in the described interactions are highly conserved and are also present in mammals (Figure S5B). Therefore, we speculate that the effect of the studied substitutions of uS7 might result in similar phenotypes in mammals.

In comparison to the py48S-eIF5N structure (Llácer et al., 2018), the electron density of eIF5-NTD was not observed in our complexes, although mass spectrometry analysis revealed some residual presence of eIF5 only in the β-globin LS48S IC. This may suggest that the presented LS48S ICs were trapped between eIF1 dissociation and eIF5-NTD binding (which would represent the structure corresponding to the intermediate state between p48S-closed and p48S-5N, according to Llácer et al., 2018). Another reason might be the weaker affinity of eIF5 to the initiation complex at this stage, as both of our complexes are purified directly from RRL without supplementation of any factors, which is overcome when the complex is assembled *in vitro* with higher molar ratios.

After the dissociation of the TC (GTP-driven binding), eIF5B is recruited to the IC at the exact binding site of ABCE1. As shown by (Wang et al., 2019), the time between binding of eIF5B and association of 60S is very short (∼0.59 s) and due to its dynamic nature, we most likely would not be able to stall the 48S-eIF5B complex using our protocol. Indeed, only a small number of MS spectra for eIF5B was recorded.

When compared to the yeast 40S ribosome-ABCE1 post-splitting complex (Heuer et al., 2017), we do not observe any large structural differences. However, NTP pocket 1 appears to be more open than NTP pocket 2 (Figures 6A and 6E, coloured frames) and it is similar to the “open state” found by X-ray crystallography (Karcher et al., 2008). Recent single-molecule-based fluorescence resonance energy transfer (smFRET) study on archeal ABCE1 showed that two NTP sites are in asymmetric dynamic equilibrium and both NTP sites can exist in different conformations (Gouridis et al., 2019). Therefore, we propose that in LS48S IC the ABCE1 is present in its asymmetric conformer, where NTP pocket 1 is in the open state, whereas pocket 2 is in the closed state. The position of ABCE1 in the IC suggests steric incompatibility with the human re-initiation factor, eIF2D (Weisser et al., 2017). Indeed, the winged helix (WH) domain of eIF2D was found to interact with the central part of h44 ribosomal RNA in the absence of ABCE1, at the exact position of the ABCE1 Fe-S cluster (I) in our LS48S IC (Figure S4B). This cluster also shows sequence similarity to the C-terminal SUI domain of eIF2D, found to be located in the re-initiation complex at the top of h44 rRNA (Figure 4A) (Weisser et al., 2017).

Regarding the relatively low abundance of eIF3 in our complexes, we believe that after the LS48S complex formation, eIF3 simply detaches from the 40S, probably during the grid preparation as has been consistently observed in structural studies of analogous complexes. The superposition of the eIF3 octamer to our previous structure of the *in vitro* reconstituted 43S PIC (des Georges et al., 2015) showed high structural similarity (RMSD 1.3 Å over all atoms). Consistent with previous study (des Georges et al., 2015), our structure reveals a sizable unassigned density at the mRNA channel exit, interacting mainly with eIF3 a and c (Figures S6B and S6C). Because of its location, it is tempting to attribute this unassigned density to the 5’UTR of mRNA, however its presence in 43S PIC (des Georges et al., 2015), which does not contain any mRNA, strongly contradicts this assignment. Thus, following our previous suggestion, we believe that this density belongs mainly to flexible segments of eIF3d. Finally, the eIF3 b-i-g module is not visible in our structure, however in the case of py48S-eIF5 IC structure (Llácer et al., 2018), it was demonstrated that these subunits relocate together to the solvent site upon start-codon recognition.

## CONCLUSIONS

Our cryo-EM structure at 3.0 Å represents the highest resolution reconstruction of a mammalian translation initiation complex till date. It refines our understanding of the architecture of late-stage IC and provides structural insights into the Kozak sequence role in canonical cap-dependent translation initiation. The data presented here demonstrate different interaction networks of the mRNA within the initiation complex based on its sequence. Furthermore, our results demonstrate that the binding of ABCE1 does not impact the conformation of the 48S IC. Finally, our structure reveals the molecular details of the mammalian eIF3 core interactions with the 40S at near-atomic resolution.

## ACKNOWLEDGEMENTS

We thank Gabor Papai and Julio Ortiz Espinoza (IGBMC, Strasbourg, France) for assistance in data acquisition and Franck Martin for providing the plasmid for β-globin mRNA. We also thank the High-Performance Computing Centre of the University of Strasbourg (funded by the Equipex Equip@Meso project) for IT support and the staff of the proteomic platform of Strasbourg-Esplanade for conducting the nanoLC-MS/MS analysis (funded by LABEX: ANR-10-LABX-0036 NETRNA). The mass spectrometry instrumentation was granted from Université de Strasbourg (IdEx 2015 Equipement mi-lourd). We thank Alan G. Hinnebusch for critical reading of the manuscript and Cameron Mackereth for numerous useful comments to it. We acknowledge Israel S. Fernández for providing us the molecular model of eIF3 core. This work was supported by the ANR grant ANR-14-ACHN-0024 @RAction program ‘‘ANR CryoEM80S’’ and the ERC-2017-STG #759120 “TransTryp” (to Y.H.).

## AUTHOR CONTRIBUTIONS

A.S. has conducted samples preparation and optimization for cryo-EM study and mass-spectrometry analysis. Y.H. and A.S. performed cryo-electron microscopy experiments. Y.H., E.G. and A.S. have interpreted the data. Y.H. and E.G. carried out data processing, structural analysis and wrote the manuscript with input from all authors. A.B. performed the atomic modelling. L.K. performed the mass spectrometry experiments. All authors read and commented the manuscript. Y.H. directed research.

## MATERIALS and METHODS

| REAGENT or RESOURCE | SOURCE | IDENTIFIER |
| --- | --- | --- |
| Chemicals, Peptides, and Recombinant Proteins |  |  |
| Guanosine 5'-[$\beta$,γ-imido]triphosphate (GMP-PNP) | Merck | Car#148892-91 |
| Protease inhibitor cocktail tablets | Roche | Cat#11873580001 |
| RNasin® Ribonuclease Inhibitors | Promega | Cat#N251B |
| Rabbit Reticulocyte Lysate nuclease-treated | Promega | Cat#L416A |
| ScriptCap™ m<sup>7</sup>G Capping System | Epicentre | Cat#SCCE0625 |
| Deposited Data |  |  |
| Structure of β-globin LS48S IC | This Paper | PDB ID: 6YAL |
| Structure of β-globin LS48S+eIF3 IC | This Paper | PDB ID: 6YAM |
| Structure of H4 LS48S IC | This Paper | PDB ID: 6YAN |
| Cryo-electron microscopy map of β-globin LS48S IC | This Paper | EMDB ID: EMD-10760 |
| Cryo-electron microscopy map of β-globin LS48S+eIF3 IC | This Paper | EMDB ID: EMD-10761 |
| Cryo-electron microscopy map of H4 LS48S IC | This Paper | EMDB ID: EMD-10762 |
| Cryo-electron microscopy map of β-globin LS48S IC + ATP | This Paper | EMDB ID: EMD-10763 |
| Cryo-electron microscopy map of β-globin LS48S+eIF3 IC + ATP | This Paper | EMDB ID: EMD-10764 |
| Recombinant DNA |  |  |
| plasmid for β-globin mRNA | (Simonetti et al. 2016) | N/A |
| plasmid for H4 mRNA | This Paper | N/A |
| Software and Algorithms |  |  |
| SCIPION | (de la Rosa-Trevín et al., 2016) | http://scipion.i2pc.es |
| Molecular Dynamic Flexible Fitting | (Trabuco et al., 2008) | https://www.ks.uiuc.edu/Research/mdff/ |
| CTFFIND4 | (Rohou and Grigorieff, 2015) | http://grigoriefflab.janelia.org/ctf |
| RELION | (Scheres, 2012) | https://www2.mrc-lmb.cam.ac.uk/groups/scheres/impact.html |
| UCSF Chimera | (Pettersen et al., 2004). | https://www.cgl.ucsf.edu/chimera/ |
| MotionCor | (Zheng et al., 2017) | https://emcore.ucsf.edu/ucsf-motioncor2 |
| PHENIX 1.9.1692<br>Phenix.ERRASER | (Adams et al., 2010)<br>(Chou et al., 2016) | http://www.phenix-online.org/ |
| Visual Molecular Dynamics | (Humphrey et al., 1996) | https://www.ks.uiuc.edu/Research/vmd/ |
| Scalable Molecular Dynamics 2 | (Phillips et al., 2005) | http://www.phenix-online.org/ |

### Lead Contact and Materials Availability

Further information and requests for reagents should be sent to the Lead Contact, Yaser Hashem.

### *In vitro* transcription and capping

Human β-globin mRNA was prepared as previously described (Simonetti et al., 2016). The template for mouse H4– 12 mRNA (375 nt; accession number X13235) was generated by PCR amplification extended on its 3′-end with a 5′-(CAA)_9_CAC-3′ tail from plasmid containing the gene synthetized by Proteogenix. The PCR product purification and in vitro transcription of mouse H4–12 mRNA were performed as described for the preparation of β-globin mRNA. The pure transcripts were capped using the ScriptCap™ m^7^G Capping System (Epicentre). Radiolabelled transcripts were obtained by substituting the GTP from the kit with [α^32^P]GTP.

### Sample preparation for Cryo-EM

β-globin 48S IC and H4 48S IC were isolated from nuclease-treated rabbit reticulocyte lysate (RRL) (Promega L4960), as previously described (Simonetti et al., 2016) with the main difference that a lower concentration of Mg^2+^ was used for ribosome complex assembly as detailed below. Prior to complexes assembly, the reaction mix in a final volume of 83 µl has been prepared by adding 21 µl of the Amino Acid Mixture (for a final concentration of 0.13 mM) and 80 units of RNasin (Promega Ref. number N2511) to 60 µl of RRL. The mix was incubated at 30°C for 5 minutes to reactivate the ribosomes. In a final volume of 157 µl, complex assembly has been obtained by adding 13 µg of mRNA, guanylyl imidodiphosphate (GMP-PNP) to final concentration of 5 mM and Mg(OAc)_2_ to keep the final concentration of free Mg^2+^ at 0.5 mM. The reaction is incubated for further 5 minutes at 30°C. β-globin and H4 mRNA assembled complexes were separate on 5%–25% linear sucrose gradient (in buffer containing 25 mM HEPES-KOH [pH 7.6], 79 mM KOAc, 0.5 mM Mg(OAc)_2_, and 1 mM DTT) by centrifugation at 36,000 rpm in a SW41Ti rotor for 4.5 h at 4°C. Moreover, we have optimized the gradient fraction collection using BioComp Piston Gradient Fractionator™ devise. The formation of translation initiation complexes has been monitored following the ribosome profile via the UV absorbance (optical density [OD] at 260 nm) and the radioactivity profile of the ^32^P-labeled globin or H4 mRNA. Fractions containing β-globin/48S IC or H4/48S IC were centrifuged at 108,000 rpm (S140AT Sorvall-Hitachi rotor) for 1 h at 4°C and the ribosomal pellet was dissolved in a buffer containing 10 mM HEPES-KOH pH 7.4, 50 mM KOAc, 10 mM NH_4_Cl, 5 mM Mg(OAc)_2_ and 2 mM DTT.

### Grids preparation and data collection parameters

The grids were prepared by applying 4 µL of each complex at ∼70 nM to 400 mesh holey carbon Quantifoil 2/2 grids (Quantifoil Micro Tools). The grids were blotted for 1.5 sec at 4°C, 100% humidity, using waiting time 30 s, and blot force 4 (Vitrobot Mark IV). The data acquisitions were performed for the β-globin•GMP-PNP and H4•GMP-PNP ICs on a Titan Krios S-FEG instrument (FEI) operated at 300 kV acceleration voltage and at a nominal underfocus of Δz = ∼ 0.5 to ∼ 3.5 μm using the CMOS Summit K2 direct electron detector 4,096 x 4,096 camera and automated data collection with SerialEM (Mastronarde, 2003) at a nominal magnification of 59,000 x. The K2 camera was used at super-resolution mode and the output movies were binned twice resulting in a pixel size of 1.1Å at the specimen level (the calibrated magnification on the 6.35 µm pixel camera is 115,455 x). The camera was setup to collect 20 frames and frames 3 to 20 were aligned. Total collected dose is ∼26 e^−^/Å^2^. In addition, Cryo-EM images of the β-globin•GMP-PNP+ATP were collected on a Polara Tecnai F30 cryo-transmission electron microscope (FEI instruments) operated at 300 keV acceleration voltage and at a nominal underfocus of Δz = ∼ 0.5 to ∼4.0 µm, using a direct electron detector CMOS (Falcon I) 4,096 x 4,096 camera calibrated at a nominal magnification of 59,000 x, resulting in a pixel size of 1.815 Å.

### Image processing

SCIPION (de la Rosa-Trevín et al., 2016) package was used for image processing and 3D reconstruction. MotionCor (Zheng et al., 2017) was used for the movie alignment of 8238 movies from the β-globin complex and 8520 movies for the H4 complex. CTFFIND4 (Rohou and Grigorieff, 2015) was used for the estimation of the contrast transfer function of an average image of the whole stack. Particles were selected in SCIPION. Approximately 1,067,000 particles were selected for the β-globin•GMP-PNP IC, 666,000 particles for the H4•GMP-PNP and 200,000 particles for the β-globin•GMP-PNP+ATP IC. RELION (Scheres, 2012) was used for particle sorting through 3D classification via SCIPION, (please refer to Figure S1 for particle sorting details for all three complexes). Selected classes were refined using RELION’s 3D autorefine and the final refined classes were then post-processed using the procedure implemented in RELION applied to the final maps for appropriate masking, B factor sharpening, and resolution validation to avoid over-fitting.

### Model building, map fitting and refinement

The four initiation complexes were modelled based on the previous initiation *Oryctolagus cuniculus* complex (PDBID: 5K0Y) (Simonetti et al., 2016) resolved at 5.8 Å. Adjustments of RNA and proteins were done using the visualization and modelling software UCSF Chimera version 1.12 (build 41623) (Pettersen et al., 2004). Sequences of modelled factors from *Oryctolagus cuniculus* were retrieved using BLAST (Altschul et al., 1990) tools in the NCBI database (NCBI Resource Coordinators, 2017) using respective template sequence described below. Templates structures were extracted from the PDB (Berman et al., 2006). ABCE1 from *Saccharomyces cerevisiae* 40S complex (PDB ID: 5LL6 chain h) (Heuer et al., 2017) was used as template to thread ABCE1 of *Oryctolagus cuniculus* in Swiss-model (Biasini et al., 2014) webservice. The core of initiation factor 3 (eIF3) composed of subunits A, C, E, F, H, K, L and M was extracted from corresponding mammalian eIF3 (PDB ID: 5A5T) (des Georges et al., 2015) and further rmodelled using (Neupane et al., 2019). Initiation factor 3D (eIF3d) from *Nasonia vitripennis* (PDB ID: 5K4B) (Lee et al., 2016) was the template to thread eIF3d of *Oryctolagus cuniculus* in Swiss-model. Eukaryotic Initiation factor 1A (eIF1A) template was extracted from *Saccharomyces cerevisiae* 48S pre-initiation complex (PDB ID: 3JAP chain i) (Llacer et al., 2015) and thread into *Oryctolagus cuniculus* in Swiss-model. The ternary complex (TC) was affined from *Oryctolagus cuniculus* initiation complex (PDB ID: 5K0Y) (Simonetti et al., 2016). Both messenger RNAs (globin and H4) were modelled using modelling tools of Chimera. Refinements were done on all four complexes in their corresponding maps. The refinement workflow followed four major steps that applied to all initiation complexes. First, a Molecular Dynamic Flexible Fitting (MDFF) (Trabuco et al., 2008) ran for 200000 steps with gscale of 1 (potential given to the density map to attract atoms in their density). The trajectories reached a plateau of RMSD curve around frame 160 for the four complexes. A minimization followed the trajectories to relax the system. MDFF ran on VMD (Humphrey et al., 1996) 1.9.2 coupled with NAMD2 (Phillips et al., 2005) v.1.3. software. Next steps of refinement required the usage of several specialized tools for RNA and proteins geometry included as modules in PHENIX (Adams et al., 2010) version 1.13-2998-000 software. Phenix.ERRASER (Chou et al., 2016) is a specialized tool for RNA refinement and Phenix.real_space_refine is specialized for proteins geometry and density fitting refinement. Finally, a last step of minimization using VMD and NAMD2 was applied. Assessment and validation of our models were done by Molprobity (Chen et al., 2010) webservice. Validation statistics are in Table S1.

### Mass spectrometry analysis and data post-processing

Protein extracts were precipitated overnight with 5 volumes of cold 0.1 M ammonium acetate in 100% methanol. Proteins were then digested with sequencing-grade trypsin (Promega, Fitchburg, MA, USA) as described previously (Khusainov et al., 2016). Each sample was further analyzed by nanoLC-MS/MS on a QExactive+ mass spectrometer coupled to an EASY-nanoLC-1000 (Thermo-Fisher Scientific, USA). Data were searched against the rabbit UniprotKB sub-database with a decoy strategy (UniprotKB release 2016-08-22, taxon 9986 *Oryctolagus cuniculus*, 23086 forward protein sequences). Peptides and proteins were identified with Mascot algorithm (version 2.5.1, Matrix Science, London, UK) and data were further imported into Proline v1.4 software (http://proline.profiproteomics.fr/). Proteins were validated on Mascot pretty rank equal to 1, and 1% FDR on both peptide spectrum matches (PSM score) and protein sets (Protein Set score). The total number of MS/MS fragmentation spectra was used to quantify each protein from three independent biological replicates (Spectral Count relative quantification). Proline was further used to align the Spectral Count values across all samples. The average of three experiments was normalized in respect to the total number of spectral counts (NSC) for all initiation factors (eIFs) (2807 for β-globin and 2126 for H4). Therefore, the H4 results were multiplied by the normalization factor of 1.32. Then, the multiplicands for different eIFs were added and the results underwent a second normalization according to the number of trypsin sites (>70% of probability) predicted by PeptideCutter provided by the ExPaSy server (https://web.expasy.org/peptide_cutter). The heat maps were generated in respect to the NSC. The coefficients of variations were calculated as standard of deviation between the normalized NSC and are presented in percentage (see Figure 1B).

### Alignments

The alignments of 8 eukaryotic species (Homo sapiens, Mus musculus, Danio rerio, Drosophila melanogaster, Ceanorhabditis elegans, Neurospora crassa, Saccharomyces cervisiae, Arabidopsis thaliana) shown in the figures, were done using Constraint-based Multiple Alignment Tool (COBALT) (Papadopoulos and Agarwala, 2007) from NCBI and visualized using BoxShade Server (ExPASy).

### Data and Code Availability

The atomic coordinates of the β-globin LS48S IC, β-globin LS48S+eIF3 IC and H4 LS48S IC have been deposited in the Protein Data Bank (PDB) under the accession numbers: 6YAL, 6YAM and 6YAN, respectively. The cryo-EM maps of β-globin LS48S IC, β-globin LS48S+eIF3 IC, H4 LS48S IC, β-globin LS48S IC + ATP and β-globin LS48S+eIF3 IC + ATP have been deposited in the Electron Microscopy Data Bank (EMDB) with the accession codes: EMD-10760, EMD-10761, EMD-10762, EMD-10763 and EMD-10764, respectively.

## SUPPLEMENTAL INFORMATION

### SUPPLEMENTAL FIGURES

**Figure S1, related to Figure 1.**
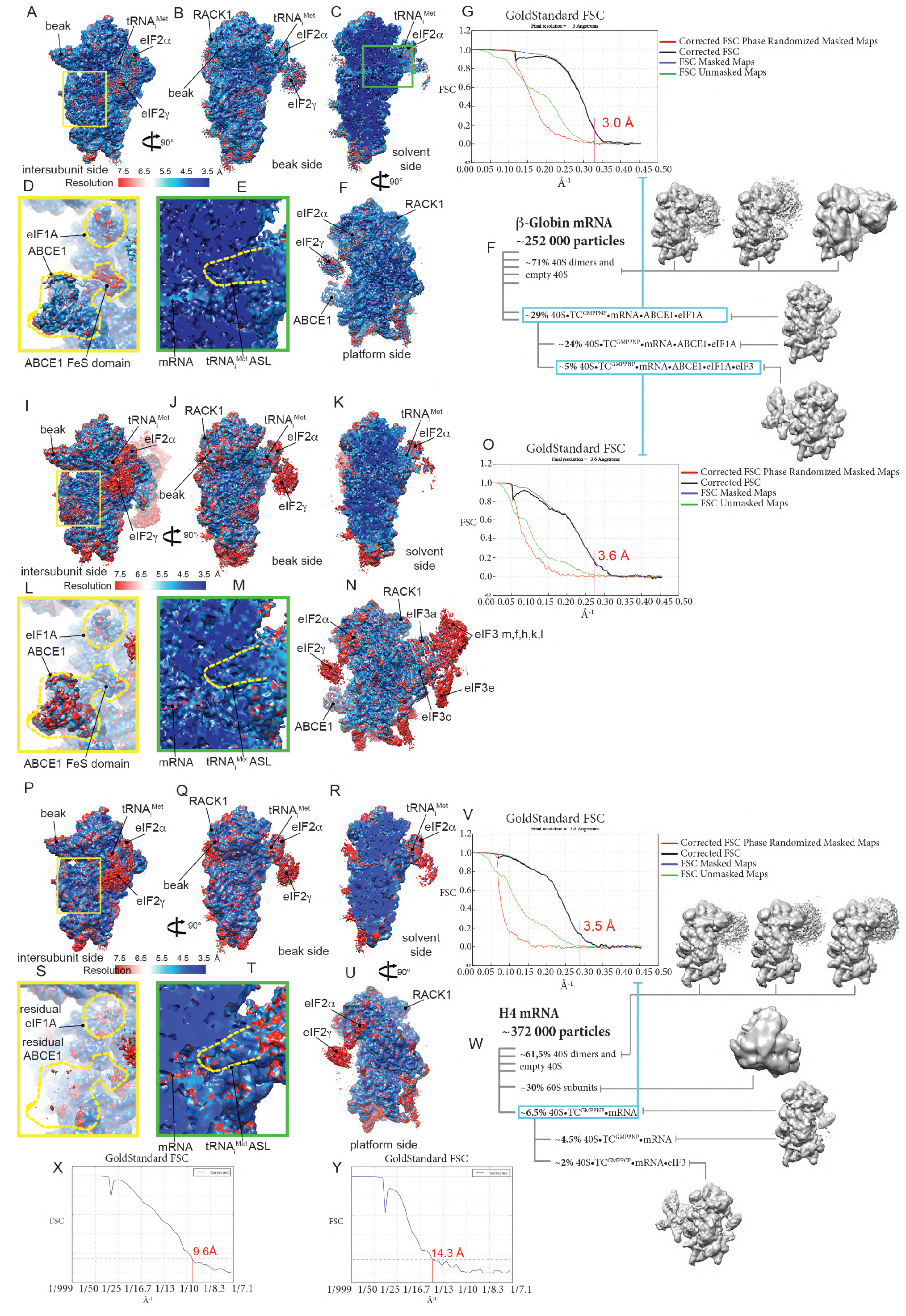
Particle classification outputs, average and local resolutions of the different β-globin and H4 LS48S ICs cryo-EM reconstructions. (A-F) Local resolution of the β-globin late-stage IC reconstruction representing a class lacking eIF3 seen from the intersubunit side (A), beak side (B), solvent side (C), platform side (F). (D-E) Insets showing local resolution of eIF1A and ABCE1 (yellow frame) and of mRNA and tRNA_i_^Met^ anticodon stem-loop (ASL) (green frame). (G) Average resolution of the reconstruction of the class without eIF3. (H) Particle classification output of the β-globin LS48S IC. (I-N) Local resolution of the β-globin LS48S IC reconstruction representing a class showing eIF3 octamer seen from intersubunit side (I), beak side (J), solvent side (K), platform side (N). (L-M) insets showing local resolution for eIF1A and ABCE1 (yellow frame) and for mRNA and tRNA_i_^Met^ ASL (green frame). (O) Average resolution of the reconstruction of the class of β-globin LS48S IC with eIF3. (P-U) Local resolution of the H4 LS48S IC reconstruction representing a class lacking eIF3 seen from the intersubunit side (P), beak side (Q), solvent side (R), platform side (U). The local resolution of eIF3 is not shown, as its average resolution is relatively low. (S-T) Insets showing local resolution of eIF1A and ABCE1 (yellow frame) and of mRNA and tRNA_i_^Met^ ASL (green frame). (V) Average resolution of the reconstruction of the class. (W) Particle classification output of the H4 LS48S IC. (X-Y) Average resolutions of the reconstruction of β-globin+ATP LS48S IC reconstructions after their particle sorting (X). Similarly to the counterpart complex with ATP, two major classes stand out, with and without eIF3 (Y).

**Figure S2, related to Figure 2 and Figure 4.**
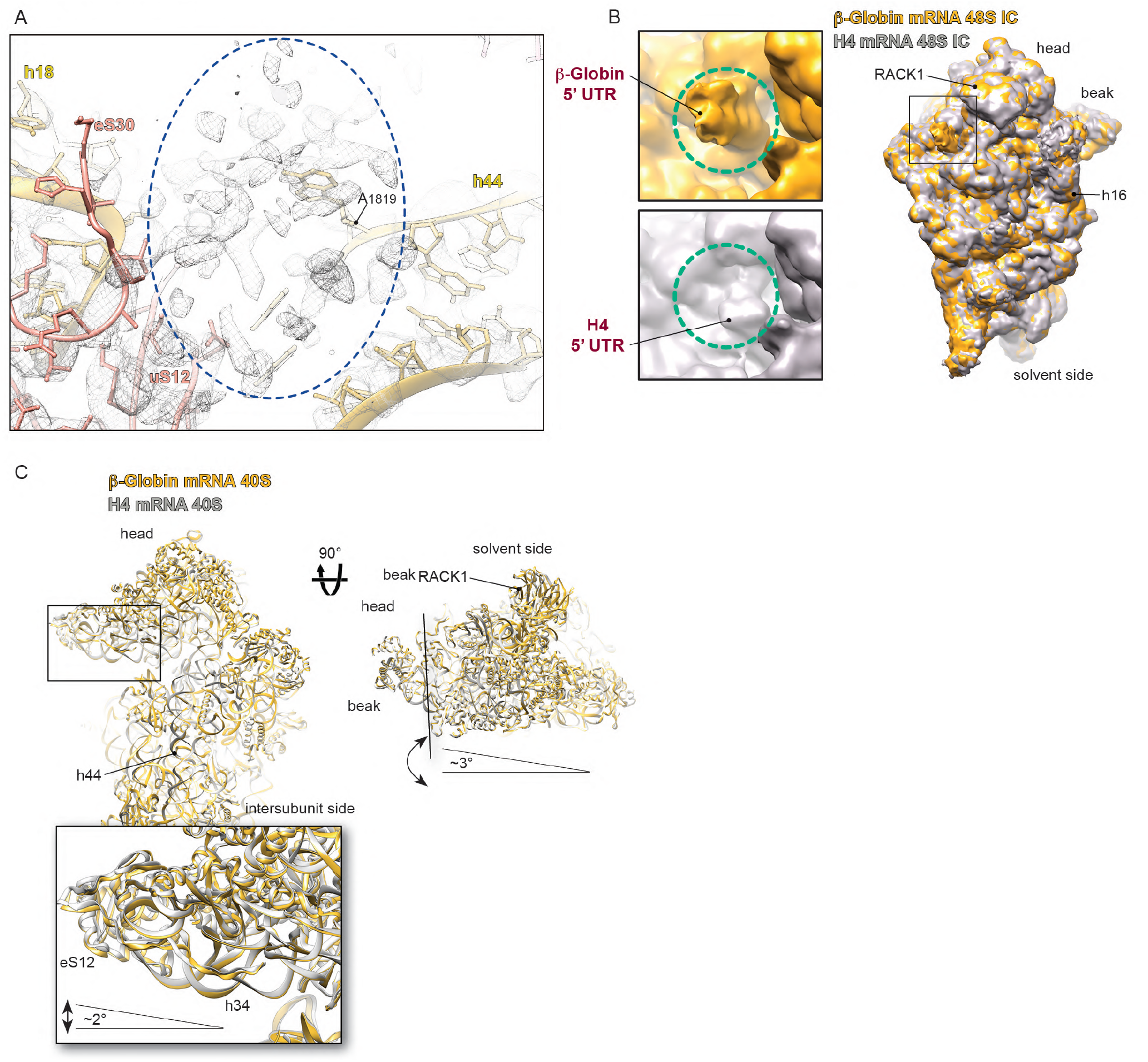
Structural differences between β-globin and H4 LS48S ICs. (A) Residual density for eIF1A in H4 LS48S IC in the A-site, highlighted by dashed blue oval. (B) Comparison of mRNA channel exit between β-globin (orange surface) and H4 (grey surface) 40S reconstructions. (C) Comparison between β-globin (gold ribbons) and H4 (grey ribbons) ICs 40S atomic models. Blowups highlight the conformational changes between both types of complexes at the mRNA channel entrance, the beak and the A, P and E –sites. Dashed coloured ovals indicate the conformational changes in several ribosomal proteins (uS3, eS30 and uS7).

**Figure S3, related to Figure 3.**
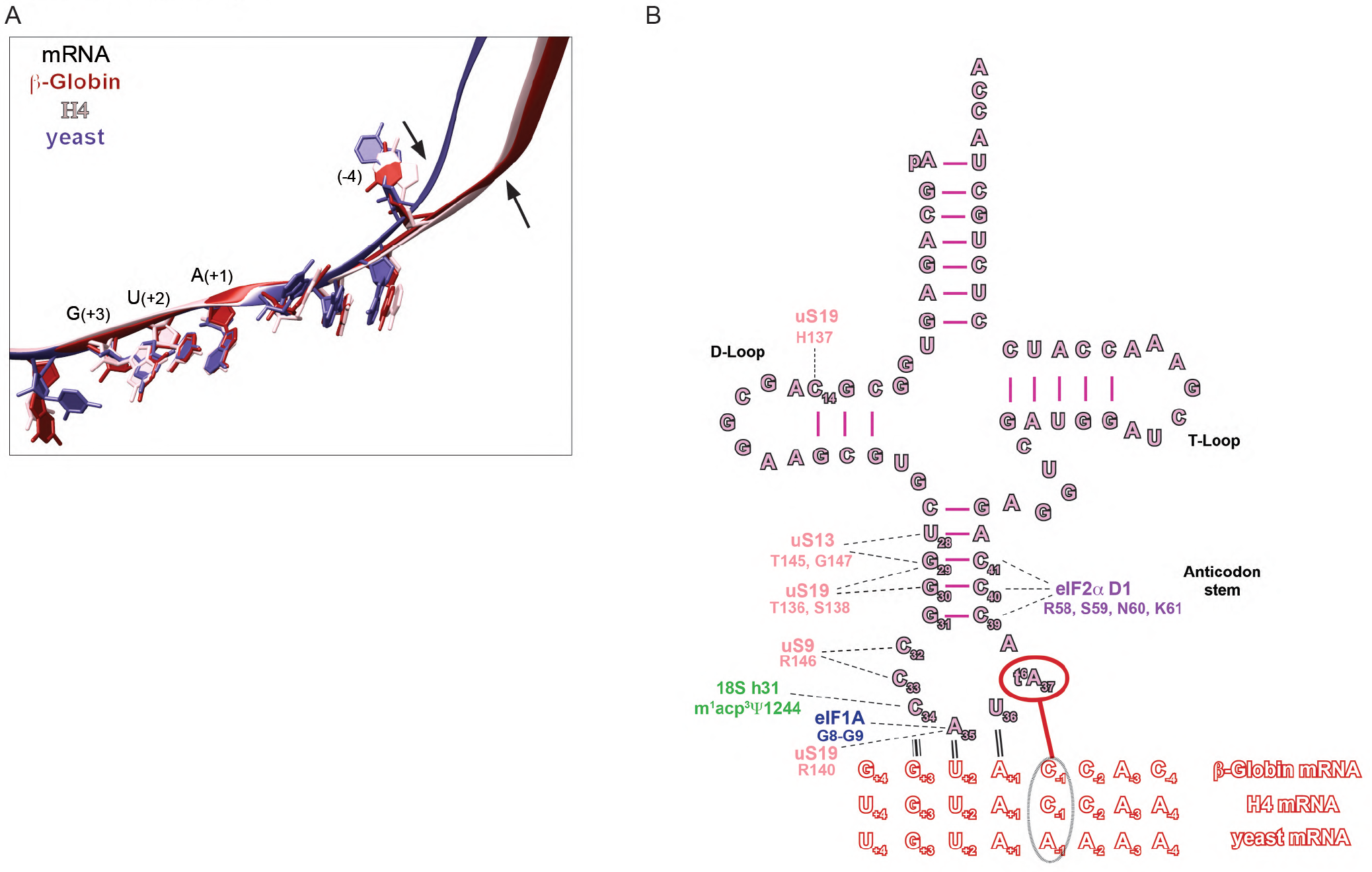
Differences between mammalian and yeast mRNA trajectories in the mRNA channel and the tRNA_i_^Met^ interactions. (A) Superimposition of yeast optimal mRNA Kozak consensus sequence in py48S-eIF5 IC structure (Llácer et al., 2018) and mammalian mRNAs showing a smoother P/E kink in the case of β-globin and H4 ICs, indicated by arrows. (B) The cloverleaf representation of the tRNA_i_^Met^ summarizing the interactions within the LS48S IC described in the Results section. The interaction of t^6^A(37) modification with (−1) mRNA position is highlighted by dashed-line oval.

**Figure S4, related to Figure 6.**
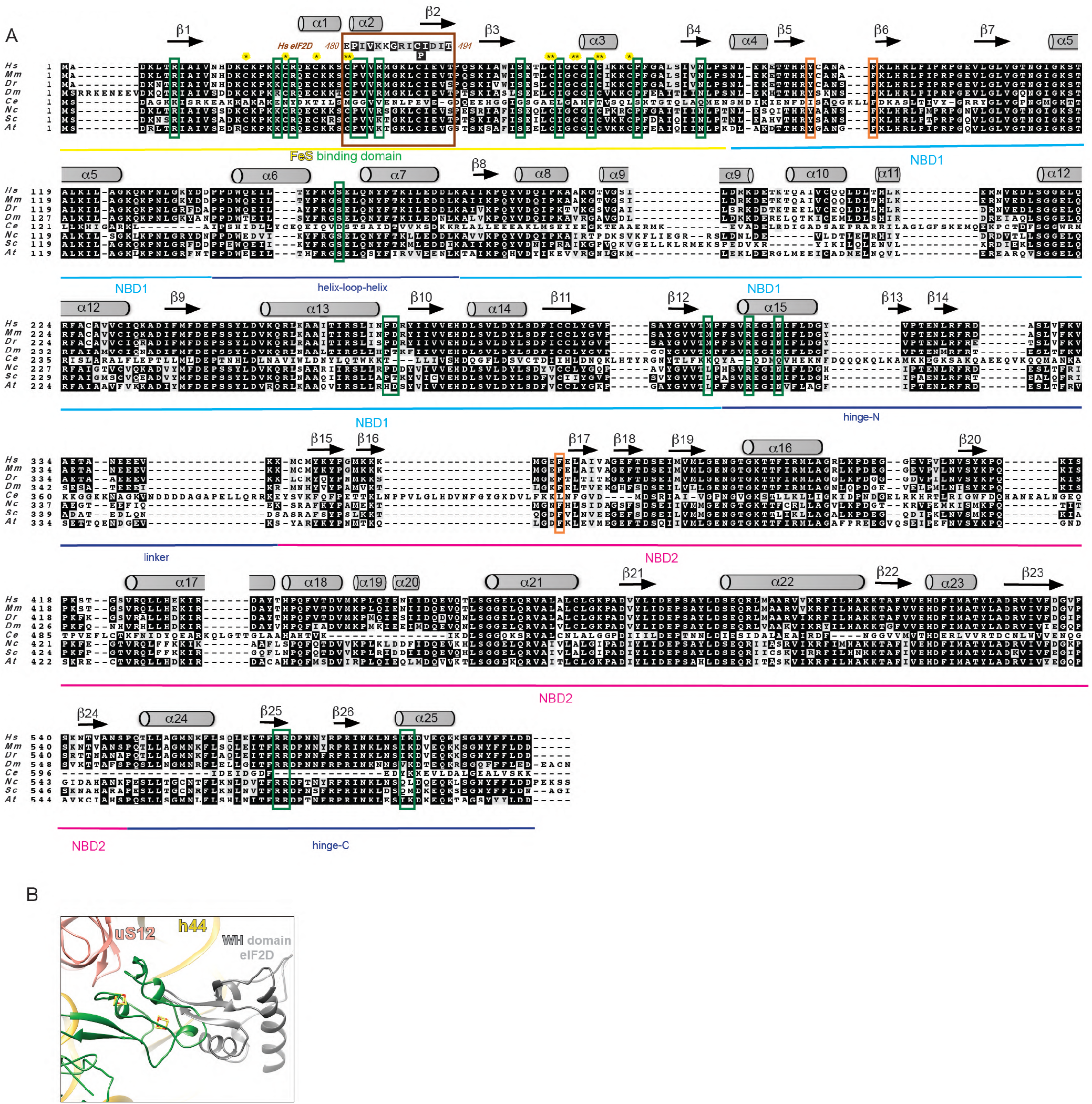
ABCE1 sequence comparison between eukaryotes and a sterical incompatibility with eIF2D. (A) Alignment of ABCE1 among eight representative eukaryotic species with secondary structure elements labelled, helices α and β-sheets (according to (Karcher et al., 2008)), and functional domains, indicated by bottom coloured lines. The residues involved in the interactions described in Results section are framed in green. In brown frame, the sequence of Fe-S cluster that showed similarity to the N-terminal part of eIF2D SUI domain (∼40% sequence identity). The coordinated cysteines of cluster (I) (*) and (II) (**) are labelled in yellow. In orange frames, the residues responsible for proper ATP/GTP binding in nucleotide binding domains (NBD1 and NBD2) pockets. The accession numbers used for the alignment: *Hs*: NP_002931.2, *Mm*: NP_056566.2, *Dr*: NP_998216.2, *Dm*: NP_648272.1 (pixie), *Ce*: NP_506192.1 ABC transporter class F, *Nc*: XP_963869.3, *Sc*: AJV19484.1, *At*: OAP04903.1). (B) Superimposition of ABCE1 in β-globin LS48S IC (in green) with eIF2D (PDB ID: 5OA3) (Gouridis et al., 2019), showing the sterical clashing of Fe-S cluster (I) with winged-helix (WH) domain of eIF2D (in grey).

**Figure S5, related to Figure 3.**
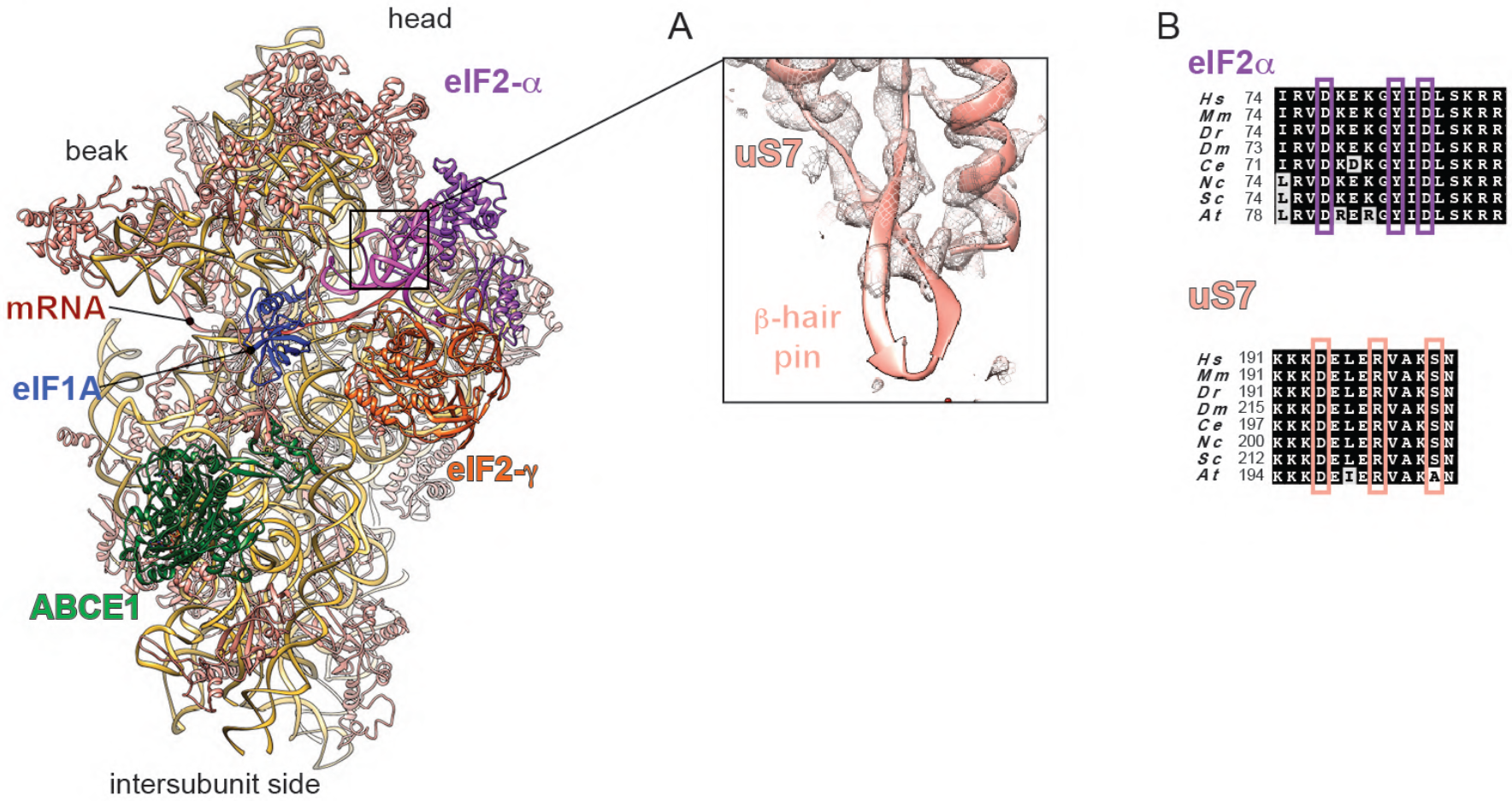
Atomic model of uS7 in mammalian LS48S IC. (A) Ribbon representation of uS7 in its electron density, showing the lack of density only in the mRNA-contacting β-hairpin, indicating the flexibility of this part of the protein. (B) Sequence conservation of the interacting residues in eIF2α and uS7, showed by (Visweswaraiah and Hinnebusch, 2017), among eukaryotic species showed in coloured frames.

**Figure S6, related to Figure 7.**
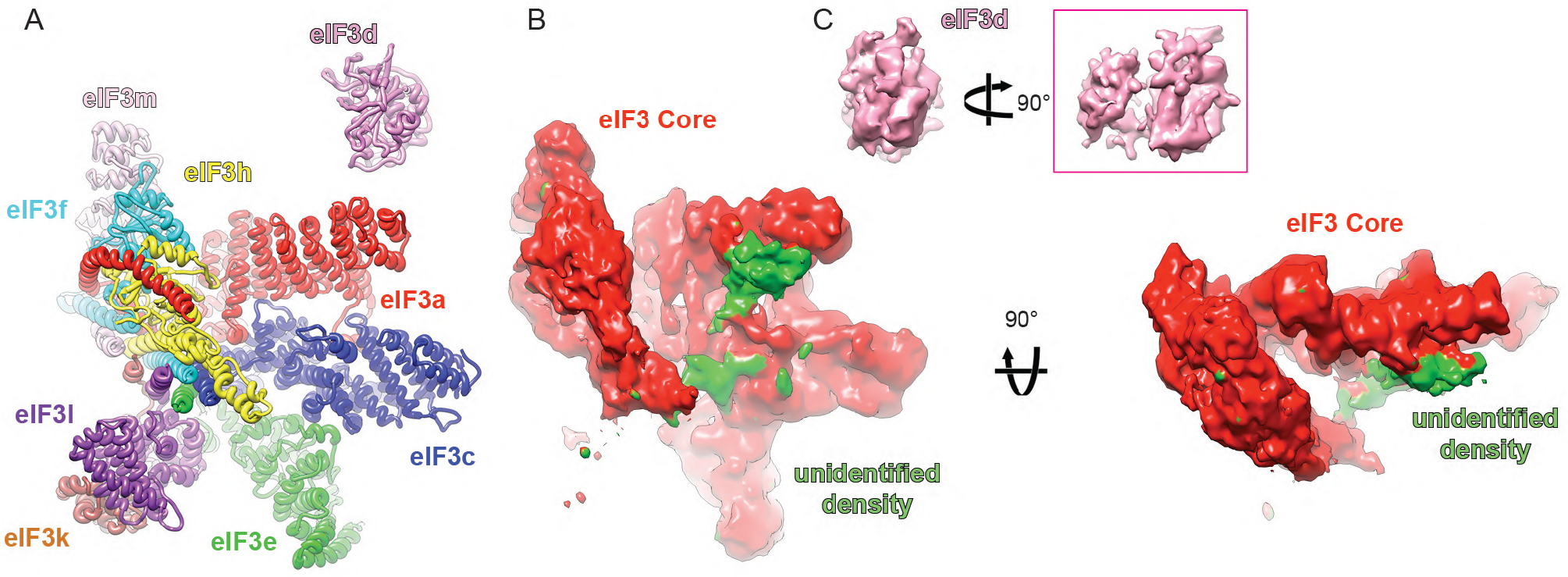
Comparison between eIF3 of ICs from near-native conditions and *in vitro* assembly. (A) Atomic model of eIF3 octamer core along with eIF3d subunit (in coloured ribbons). (B) Segmented cryo-EM densities of eIF3 octamer core along with eIF3d (filtered to 8 Å), showing unidentified density (green surface) in contact with eIF3 a and c subunits. (C) Fitting of eIF3d partial crystal structure into its cryo-EM density.

### SUPPLEMENTAL TABLES

**Table S1, related to Figure 1.**
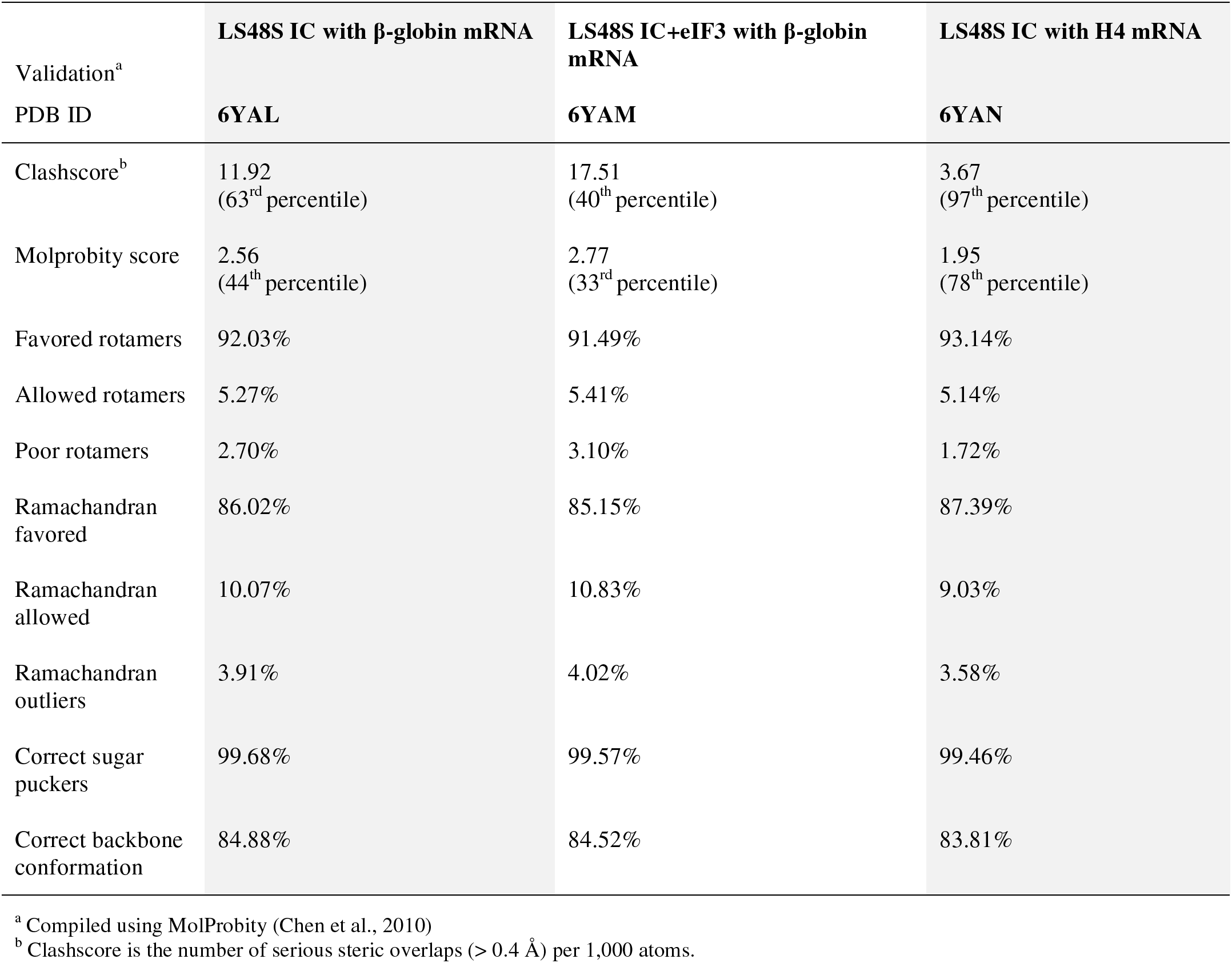
Refinement and validation statistics for the three LS48S IC structures from *O.cuniculus*.

**Table S2, related to Figures 2-7.**
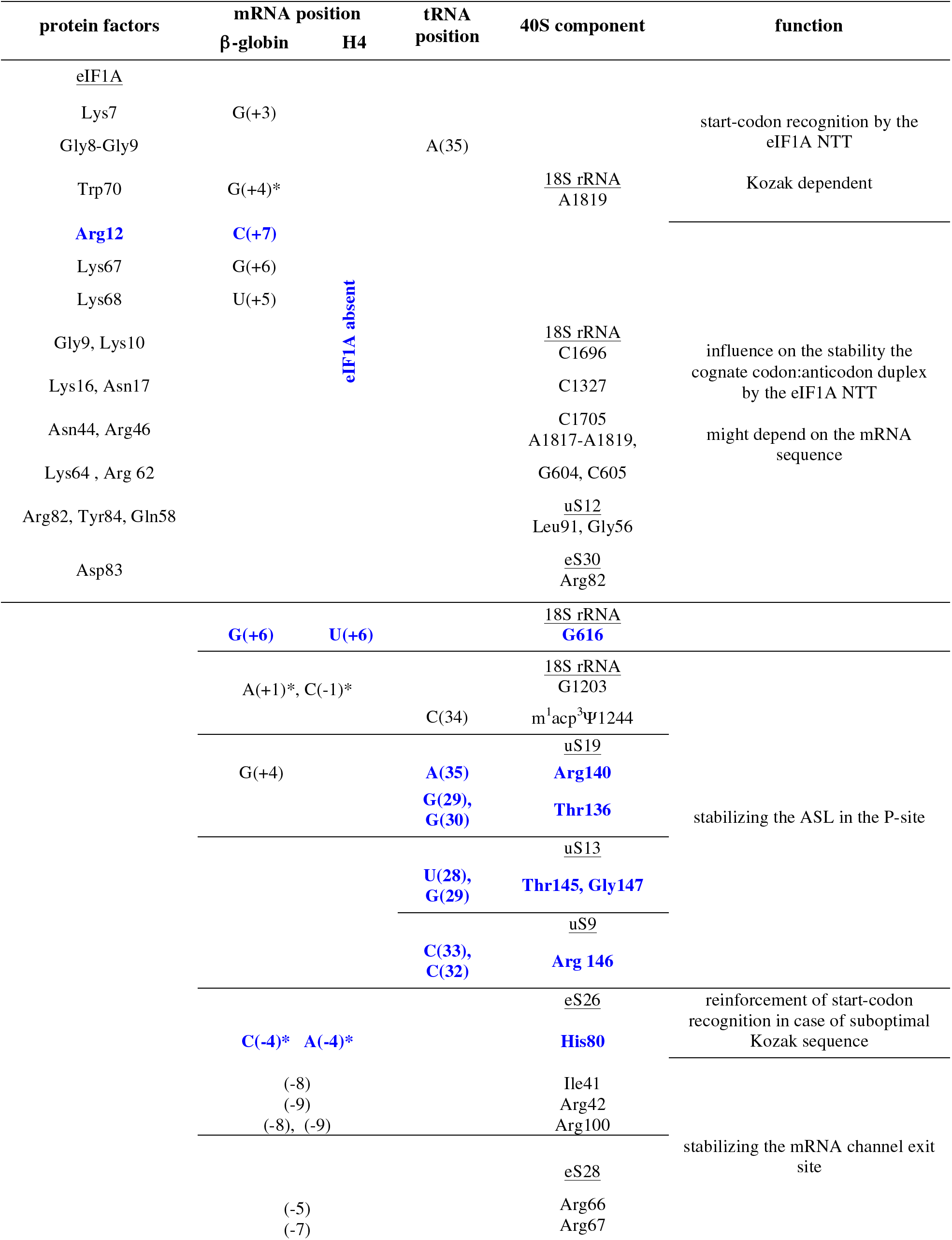
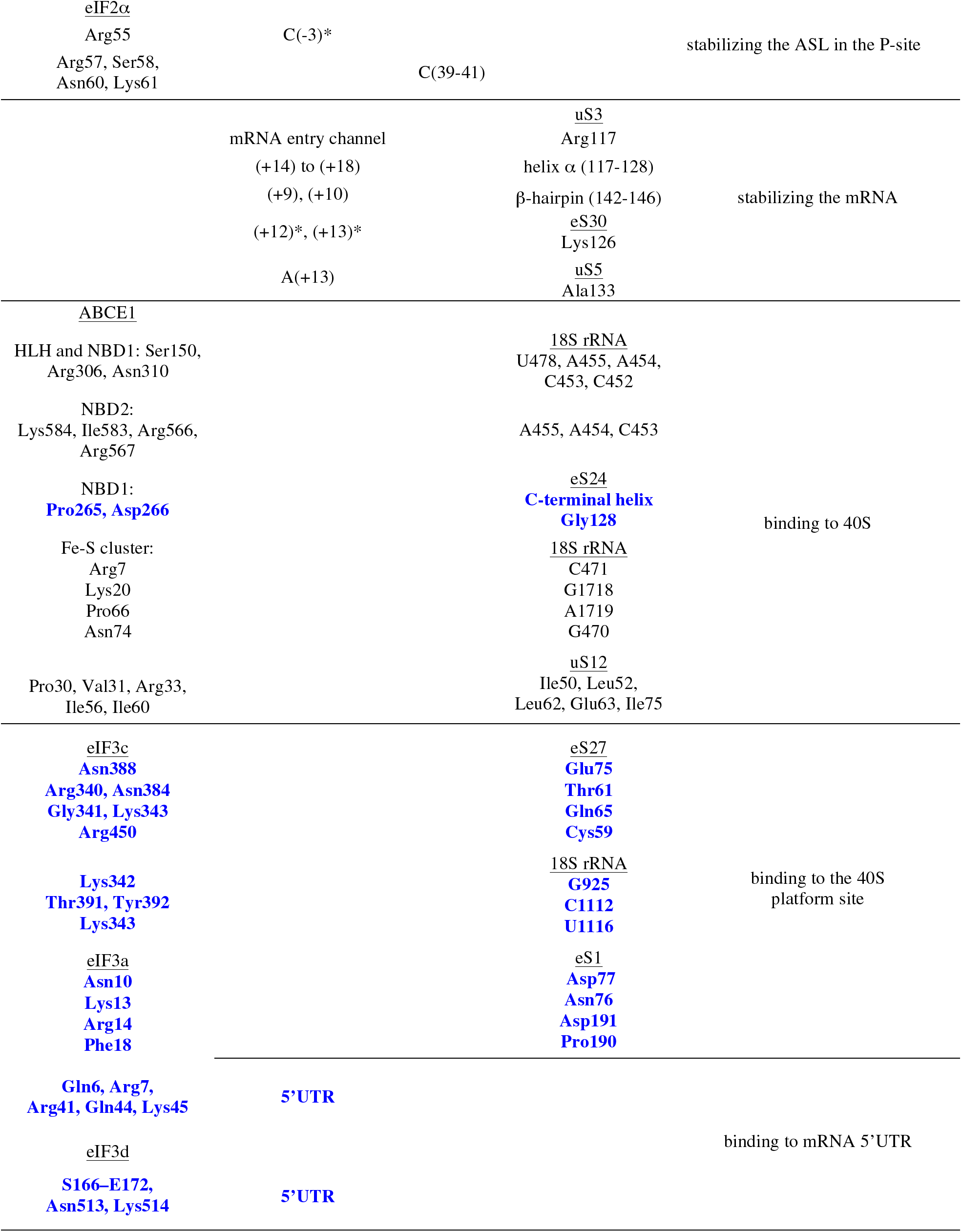
Summary of the interaction found in the β-globin LS48S and H4 LS48S ICs, described in this paper. The contacts found in this work are highlighted in blue. Stars indicate the interactions with the nucleotide bases.

